# HiDiver: A Suite of Methods to Merge Magnetic Resonance Histology, Light Sheet Microscopy, and Complete Brain Delineations

**DOI:** 10.1101/2022.02.10.479607

**Authors:** G. Allan Johnson, Yuqi Tian, Gary P. Cofer, James C. Cook, James C. Gee, Adam Hall, Kathryn Hornburg, Yi Qi, Fang-Cheng Yeh, Nian Wang, Leonard E. White, Robert W. Williams

## Abstract

We have developed new imaging and computational workflows to produce accurately aligned multimodal 3D images of the mouse brain that exploit high resolution magnetic resonance histology (MRH) and light sheet microscopy (LSM) with fully rendered 3D reference delineations of brain structures. The suite of methods starts with the acquisition of geometrically accurate (in-skull) brain MRIs using multi-gradient echo (MGRE) and new diffusion tensor imaging (DTI) at an isotropic spatial resolution of 15 μm. Whole brain connectomes are generated using over 100 diffusion weighted images acquired with gradients at uniformly spaced angles. Track density images are generated at a super-resolution of 5 μm. Brains are dissected from the cranium, cleared with SHIELD, stained by immunohistochemistry, and imaged by LSM at 1.8 μm/pixel. LSM channels are registered into the reference MRH space along with the Allen Brain Atlas (ABA) Common Coordinate Framework version 3 (CCFv3). The result is a **hi**gh-**d**imensional **i**ntegrated **v**olum**e** with **r**egistration (**HiDiver**) that has a global alignment accuracy of 10–50 μm. HiDiver enables 3D quantitative and global analyses of cells, circuits, connectomes, and CNS regions of interest (ROIs). Throughput is sufficiently high that HiDiver is now being used in comprehensive quantitative studies of the impact of gene variants and aging on rodent brain cytoarchitecture.

## INTRODUCTION

Magnetic resonance imaging (MRI) of the human brain has become a mainstay of both clinical care and basic neuroscience research since Lauterbur’s catalytic studies in the 1970s [1]. Resolution, contrast, and imaging strategies have improved at a rapid rate. However, resolution is ultimately limited in humans by movement artifacts, sensitivity, and short scan durations. Voxels are typically about 1 mm on edge. As a consequence, structural and functional MRI methods in human are unable to resolve key cellular and synaptic levels that are essential to probe and parse human cognition and behavior at fine-grained anatomical and physiological levels. But these studies have been enormously rewarding, especially in human clinical studies, because throughput is so high, and sample sizes can include hundreds, and even thousands of subjects.

As a result of this high throughput there is now a burgeoning new field that combines MRI methods with classical genetics and genome-wide association studies (GWAS)—for example the Human Connectome Projects [2] [3]. These joint MRI and genetic studies are now sufficiently well powered to detect DNA variants and haplotypes that are causally linked to both normal differences in brain architecture and differential susceptibility and resilience to neurological diseases. Examples include ENIGMA [4], the UK Biobank [5], and the Alzheimer’s Disease Neuroimaging Initiative [6, 7]. In comparison, MRI studies using small mammals—rats and mice in particular—have taken an orthogonal approach in which the major focus has been to achieve ever higher spatial resolution, rather than higher throughput This has involved new high-field MRI acquisition strategies from postmortem specimens, i.e., magnetic resonance histology (MRH) [8] [9] [10] [11]. We have reached the point at which an entire mouse brain can be imaged at resolutions of better than 25 μm [12], and as shown here, at a DTI super-resolution of 5 μm. These murine MRH data sets can be merged with whole-brain LSM immunohistochemistry (IHC) to provide up to two orders of magnitude higher spatial resolution. The result is that whole brains of mice can now be imaged in concert across many modalities—MRH, IHC, *in situ* hybridization, and using single cell labeling techniques.

The method we introduce here—HiDiver—exploits the inherent geometric accuracy and multiple contrasts of MRH and DTI as a veridical spatial foundation for registration of all other modalities. The result is unprecedented geometric accuracy across all modalities of imaging and across the entire brain. The minimization of distortion is critical when merging multiple data sets from single brains. Without precise alignment the high nominal resolution of LSM is far less useful, and downstream quantitative metrics become blurred or unreliable.

Achieving registration accuracy for LSM can come at a cost. Rodent brain MRH and LSM studies typically have modest sample sizes—often less than 20 cases per study and often include only one or two genotypes, ages, or treatment groups. The bottleneck is long acquisition times for both MRH and LSM—hours, days, and even weeks. There is an equally serious computational constraint in processing, aligning, and merging terabyte 3D images. These problems have thwarted high throughput rodent neurogenetics of the types heralded in humans by ENIGMA [4], ADNI [7] and UK Biobank [5].

Here we have developed and proofed the HiDiver suite of methods to accurately register MRH, LSM, with comprehensive 3D volumetric labeling of 180 ROIs per hemisphere. We demonstrate that HiDiver generalizes to different sexes, ages, and genotypes with an accuracy approaching the single cell level. Both imaging and computational workflows have a throughput of hundreds of specimens per year. This enables systematic global analyses and genetic dissection of variation in CNS architecture as a function of genotype, age, sex, environment, and treatment at both regional and cellular levels.

## RESULTS

We address core technical challenges related to methods of imaging and registering MRH with LSM for brain data sets and with 3D labels for hundreds of brain ROIs. The main challenges are: **1.** acquisition of MRH at microscopic resolution with adequate throughput; **2.** accurate registration of geometrically distorted LSM data with MRH data, and **3.** workflows to convert multimodal data sets into a common volumetrically rendered reference space. In previous studies we have demonstrated the utility of MRH in neurogenetics at moderate throughput—12 strains with two replicates each [13]. For that work we registered MRH data into the INCF Walxholm reference space [10] and generated useful volumetric data for 35 ROIs. In the last decade we have greatly increased MRH resolution, contrast, and content using diffusion tensor imaging [14] [12]. DTI allows track density imaging (TDI) and connectomics [15] [16]—as shown here at resolution down to ~ 5 μm.

**Table 1** summarizes three sets of experiments used to develop and validate HiDiver:

1. Young adult C57BL/6 cases studied at 90 days of age—an age close to that of most data generated by GENSAT [17], the ABA, and the BRAIN Initiative [18] [19]. All cases were scanned using state-of-the-art MRH for the creation of a full volume reference atlas of brain ROIs.
2. A second set of cases was used to explore the impact of spatial resolution and numbers of scan angles on scalar metrics with the goal of improving throughput and testing LSM registration accuracy. For this purpose, we generated super-resolution TDI at 5 μm resolution. Finally, we registered LSM into the C57BL/6J MRH reference volume. For this work we used a congenic C57BL/6J transgenic line generated by Sanes and colleagues (B6.Cg-Tg(*Thy1*-YFP)HJrs/J [20] [21]) that expresses yellow fluorescent protein intensely in subsets of large neurons and their axons, particularly pyramidal cells in cortical layer 5 and in hippocampus.
3. The final experiment tested the robustness and replicability of HiDiver registration in the face of significant variation in genotype and age of specimens. For this work we used BXD89 mice, an inbred strain generated by crossing of C57BL/6J to DBA/2J [22]. Its genome has been fully sequenced [23](Ashbrook et al. bioRxiv, in progress), and differs from the reference genome at ~2.87million loci—87% SNPs and 13% other variant classes (DG Ashbrook, personal communication). One of the BXD89s was a young adult at 111 days; the other was a very old adult at 687 days. Mean life expectancy of this strain is ~700 days [24].

**Table 1.**
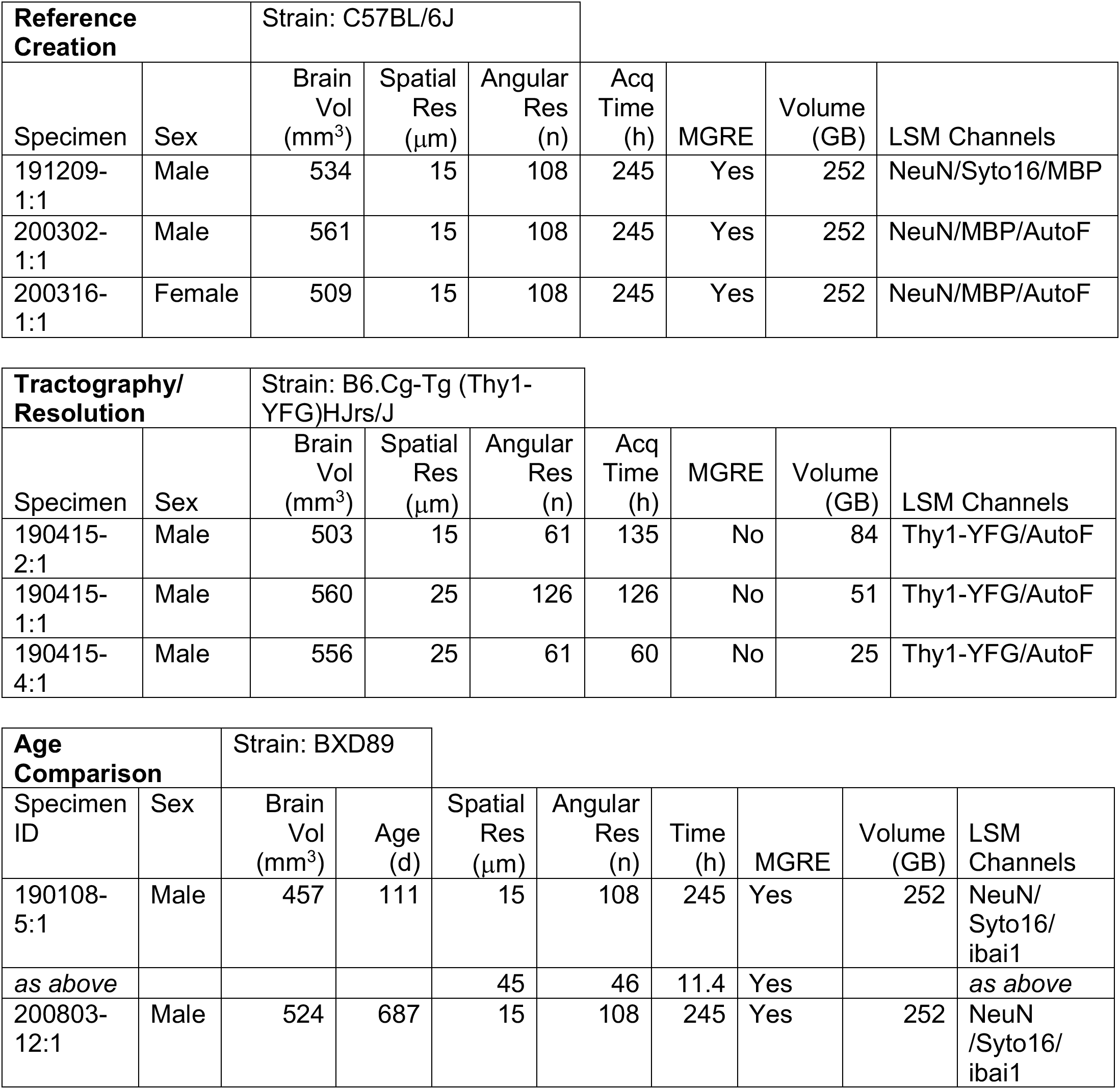
Details of specimens and acquisition protocols for three different HiDiver experiments

### Experiment 1. Reference Framework Creation

MRH resolves histological features without distortion, shrinkage, or the registration challenges that complicate most histological processing that make quantitative analysis and stereology challenging [25]. **Figure 1** provides an overview of the MRH modalities of multigradient echo image (MGRE) and scalar images from the DTI used to differentiate CNS boundaries as typically defined by conventional histological methods such as Nissl (**Fig 1A**). The average MGRE (**Fig 1B**) combines four separate echoes of a 4D MGRE volume. Scalar diffusion images are derived from the DTI [26] and GQI algorithms [27] using the 4D diffusion volume. We assembled a large (252 GB) volume by registering 108 3D diffusion weighted images each acquired with the diffusion gradient at a unique angle as well as using 13 baseline (b0) volumes (*Methods*). The resulting scalar images yield unique contrast that is related to the tissue cytoarchitecture, e.g., axon densities and myelin content. These MRH base data sets on which HiDiver is reliant consist of 12 complementary views of cyto- and myeloarchitecture, five of which are shown in **figure 1**. **Table S2** summarizes all the modalities in the standard HiDiver/MRH protocol. Blood vessels are most prominent in the MGRE and AD images (**Figs 1B,C**), whereas mean diffusivity highlights hippocampal layers particularly well (MD, **Figs 1D,d**). In marked contrast, the fractional anisotropy (FA, **Figs 1E,e**) strongly accentuates white matter tracts and those regions in which dendritic arbors have a strong common orientation, such as stratum radiatum and the molecular layer of the dentate gyrus. FA exhibits some rough similarity to acetylcholinesterase and cytochrome oxidase staining—a feature that is particularly notable in tangential sections through the barrel field (**Fig 1E arrow** and for tangential view of barrel cortex using the FA modality see **Fig S1**).

**Figure 1.**
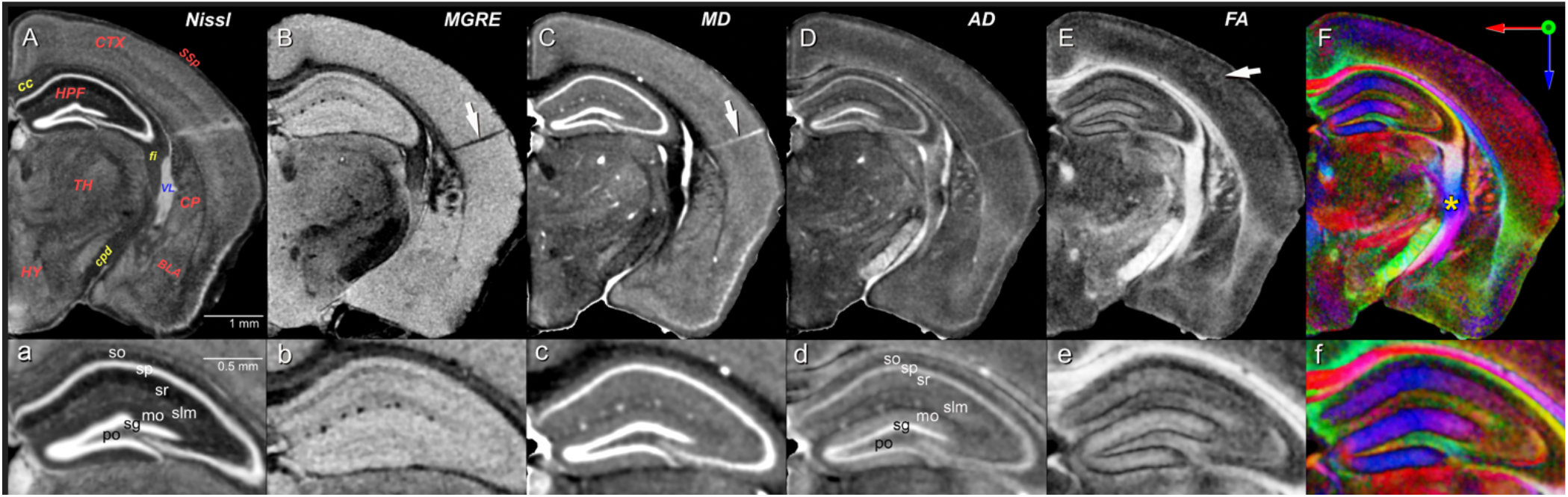
Overview of magnetic resonance histology contrast modalities at a plane equivalent to level 74 of the ABA coronal series (https://tinyurl.com/CCFv3-Level74). **A.** Reference atlas image taken from the ABA of an inverted contrast Nissl-stained section subsampled at a nominal 10 μm resolution. For orientation, several regions and tracts are labeled in **red** and **yellow**: cortex (**CTX**), corpus callosum (**cc**), primary somatosensory area (**SSp**), hippocampus (**HPF**), fimbria (**fi**), thalamus (**TH**), hypothalamus (**HY**), basolateral amygdala (**BLA**), and lateral ventricle (**V**L, in **blue**). All MRH images are closely aligned virtual sections taken from a 90-day-old C57BL/6J male (Table 1, specimen 200302-1:1). **B.** Average **MGRE** with dark blood vessels (**arrow** indicates a perforating artery) and darker heavily myelinated tracts (e.g. **cc**, **cpd**, and **fi** in **A** and **B**); **C.** Mean diffusivity (M**D**) with light regions defining both cell-dense regions such as the granule cell layer (Fig 1d, **sg**) dentate gyrus, and blood vessels (**arrow**) and ventricle. **D**. Axial diffusivity (A**D**) is similar to **C**, but differentiates additional hippocampal layers. **E**. Fractional anisotropy (**FA**) defines finer details of neurite orientation—white in the heavily myelinated corpus callosum and corticofugal projections, and the margins of barrels in somatosensory cortex (**arrow**). The entire barrel field cortex is revealed in an en-face tangential plane in **Fig S1**. **F.** ColorFA provides additional visual dissection of fiber tracts based on their orientation (top right **inset**). The yellow asterisk marks a complex of intersecting tract around the internal capsule (details with ABA overlay of tracts in **Fig S2**). Note that the corpus callosum (**red**) is distinct from both the cingulum (**green** above) and the dorsal hippocampal commissure (**green** below). **Lower row (a–f)** are corresponding 2X magnifications of the hippocampus that illustrate MRH resolution of layers. Abbreviations—**so**: stratum oriens; **sp**: pyramidal cell layer; **sr**: stratum radiatum; **slm**: stratum lacunosum-moleculare; **mo** molecular layer of the dentate gyrus; **sg**: granule cell layer of the dentate gyrus; **po**: polymorphic layer of the dentate gyrus.

The utility of color FA is emphasized by the complex intersection of tracts in the area around the internal capsule—the yellow asterisk in **figure 1F**. The image intensity represents the magnitude of diffusion anisotropy whereas color encodes the primary axis of diffusion (**Figs 1F**). The intersecting tracts in this region—the fimbria, the stria terminalis, the cerebral peducle, the internal capsule, and the optic tract—are exceedingly difficult to resolve by conventional histology. However, all these tracts can be dissected unambiguously in the colorFA. A comparative view of this region with the ABA tract demarcations is given in **figure S2**. By combining and contrasting multiple aligned MRH modalities one can computationally enhance the definition of nuclear and tract boundaries and often resolve structures and gradients that are hard to detect using standard light or even serial-section electron microscopy.

#### Definition of ROIs by transfer of ABA CCFv3 labels

The ABA CCFv3 defines ROIs for a total of 461 structures [28]. We registered all of these independently into each hemisphere of the C57BL/6J in **figure 1** using ANTs [29](**Methods**). More than half of all ROIs were so small that they often could not be reliably registered even within a single strain, sex, and age that matches that of the ABA. Registering across genotypes was even more challenging, and this compromises the accuracy with which we were able to estimate volumes, cell densities, tract DTI metrics, and total cell counts per ROI. With few exceptions almost all ROI volumes with a left-right coefficients of error (CE) greater than 0.05 are generated by poor technical accuracy rather that fluctuating asymmetry in brain structure (for detailed discussion see [30] [31]). We therefore opted to generate a reduced but comprehensive subset of ROIs that combine smaller adjacent sub-volumes. We refer to this subset as the first reduced version of CCFv3 or **r1CCFv3**. It consists of 179 bilateral ROIs and provides a full and isotropic parcellation of the brain (**Fig 2**, **Methods**, **Table S2**).

**Figure 2.**
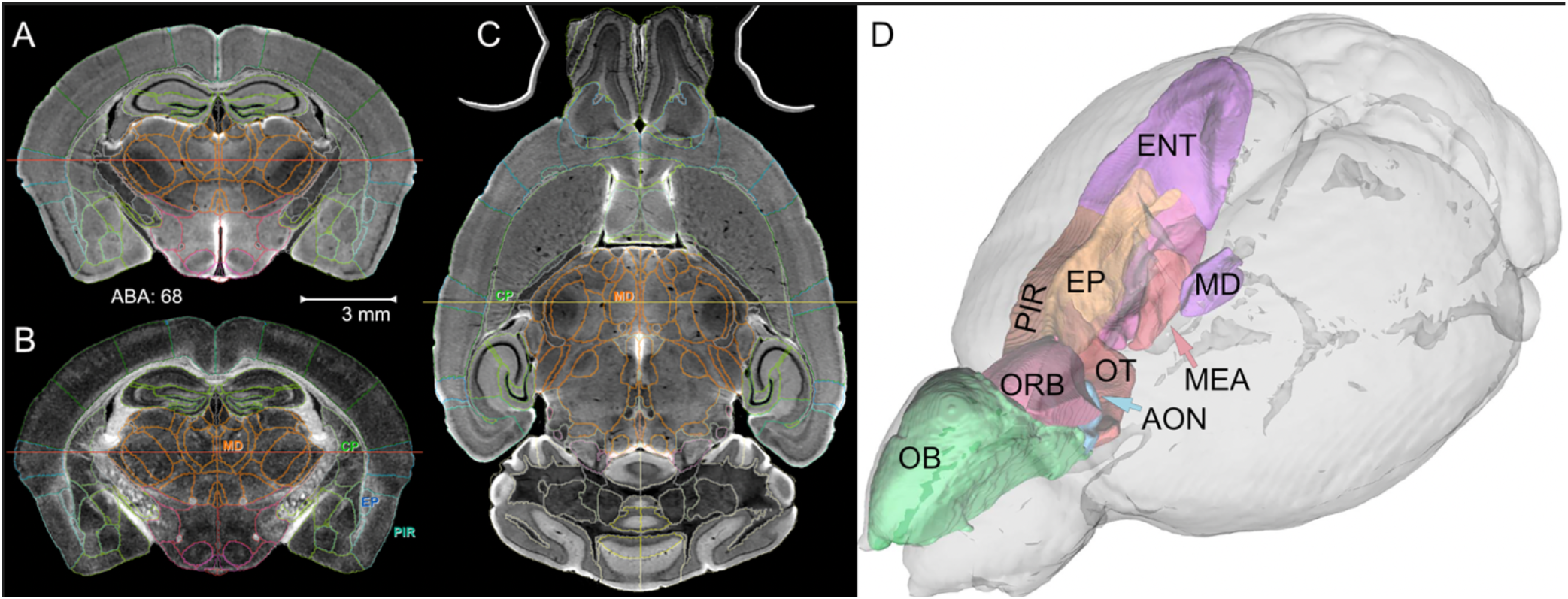
Full brain volumetric rendering of r1CCFv3 with MRH. All lines and boundaries that define ROIs use the ABA conventions (see ABA level **68** at https://tinyurl.com/CCFv3-Level68). **A.** The DWI modality in a coronal plane with ROI demarcations. **B.** The FA modality at the same level with four labeled ROIs: the mediodorsal nucleus of the thalamus (**MD**), the caudoputamen (**CP**), the endopiriform nucleus (**EP**), and the piriform area (**PIR**). **C.** Axial DWI of a horizontal section with corresponding borders and two ROIs labeled in common with **B**. The **red** lines in **A** and **B**, and the **yellow** line in **C** define orthogonal images. **D.** Full 3D delineations of major components of the olfactory system displayed with DSI Studio. Abbreviations—**OB**: olfactory bulb; **AON:** anterior olfactory nucleus; **ENT**: entorhinal area; **MEA**: medial amygdalar nucleus; **COAz**: cortical amygdalar zone; **OT**: olfactory tubule. Other abbreviations as in **B**. **Technical note:** The full set of 3D labels was mapped onto all 12 modalities summarized in Table S2 providing a rich feature set to drive subsequent registration to new strains.

The volume parcellation of r1CCFv3 was refined and corrected by two neuroanatomists (LEW, RWW). The end product was a set ROIs that can be embedded reliably and bilaterally in all MRH data sets, regardless of sex, age, and genotype, and with an acceptably low mean CE (<0.05) without extensive manual curation.

### Experiment 2. Lightsheet and MRH: HiDiver Registration

There are fundamental and often complementary constraints of different imaging modalities. MRH provides excellent geometric fidelity, multiple tightly registered contrasts (**Fig 1**), and the practicality of conducting whole brain morphometry of *in situ* volumes, ROI histology, and tractography. The major benefits of LSM are high spatial resolution—potentially down to 0.1 μm—, and the application of innumerable molecular and cellular labeling and staining protocols. The main drawback is poor geometric fidelity and problematic penetration of reagents. HiDiver makes it possible to combine the strength of both methods. **Figure 3** shows an original LSM data set before (**red**) and after (**green**) registration to the MRH data. Volumetric swelling using SHILED is over 60% relative to the MRH reference volumes (**Table 1**), and this expansion is not uniform. For example, swelling in the olfactory bulb is nearly 100%.

**Figure 3.**
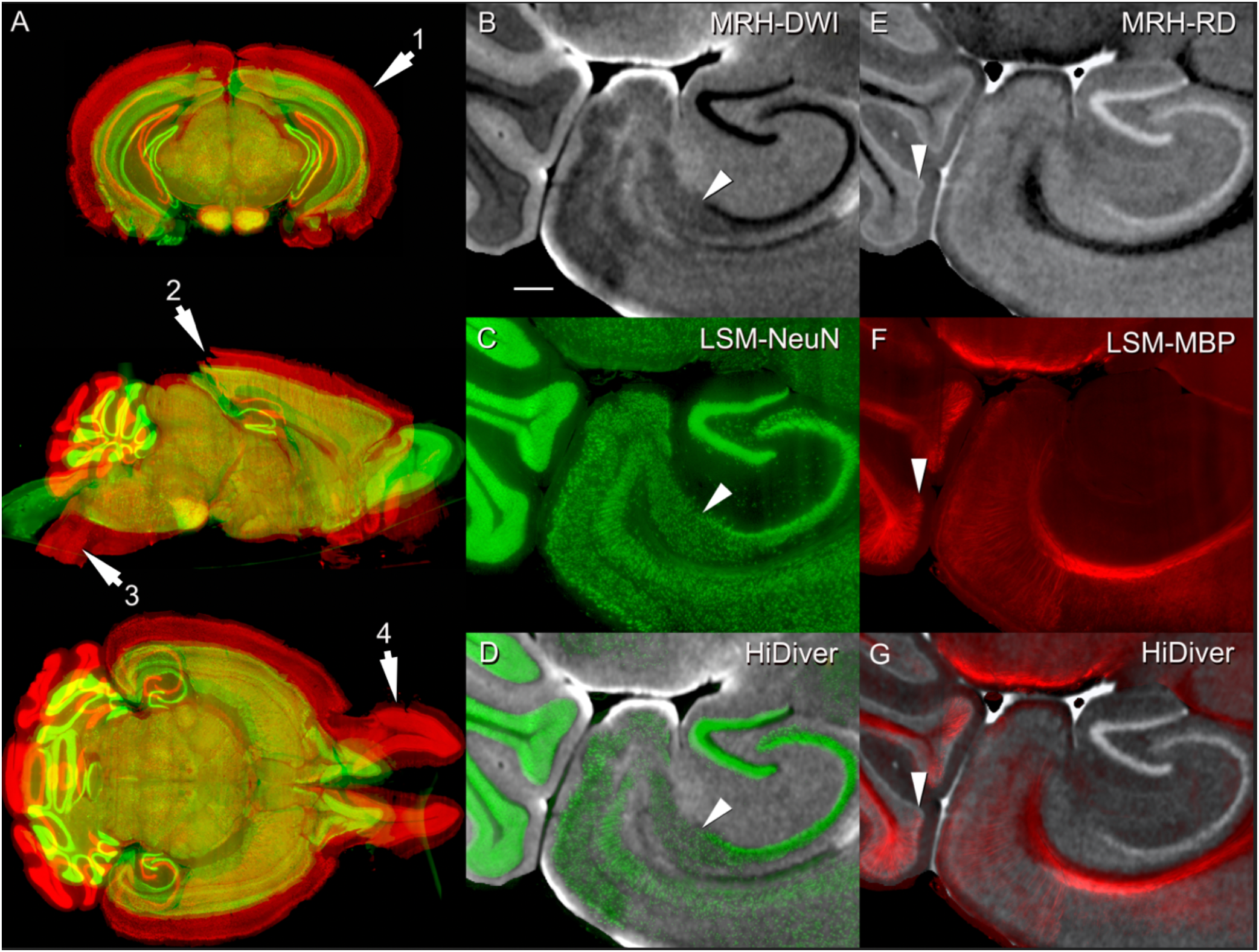
HiDiver integration of MRH and LSM channels. Swelling and distortion of LSM samples were corrected by registering to the MRH reference volume of the same specimen. Panel **A** flags four types of registration problems in the original LSM data—before (**red**) and after (**green**) correction with specimen 200316-1:1. **Arrow 1** highlights variable expansion during LSM processing that can range up to 100% for the olftactory bulbs, but averages about ~60% globally in this particular case. **Arrow 2** flags tears in visual cortex introduced during dissection or processing. **Arrow 3** marks an exaggerated flexure of the brainstem, while **arrow 4** indicates spread of the olfactory bulbs. All three LSM channels were brought back into alignment with MRH by diffeomorphic registration (**Methods**) with an accuracy that is almost always better than 50 μm. The corrected volume shown in **green** has been modified in all dimensions and consequently there is no longer a 1-to-1 correspondence between red and green versions in single planes. Panels **B, C,** and **D** (specimen 191209-1:1) provide a qualitative overview of HiDiver alignment where **B** is DWI, **C** is NeuN, and **D** is the HiDiver product. The arrowheads in these overlays point to the CA1 to subiculum boundary. **E**, **F**, and **G** are a similar trio that point to the Purkinje cell layer (a single rank of Purkinje cells and Bergmann glia) in cerebellar cortex. The MBP label extends into, and just above this layer. Image overlays were produced using Imaris. Scale bar in **B** is 200 μm.

**Figure 3** highlights two key elements relevant to the resolution and alignment of HiDiver. First, the resolution of MRH is already at the threshold of resolving single large cell bodies and fascicles of axons—15 to 25 μm. HiDiver alignment is extraordinary given the LSM distortions. Trios of oblique arrowheads in **figure 3B, C**, and **D** point out the tight correspondence of the CA1–subiculum boundary—an accuracy of about 20 to 40 μm. A similar trio of images (**Figs 3E, F**, and **G**) demonstrates the melding of MRH-RD and LSM-MBP labeling of the Purkinje cell layer. It is worth pointing out, that like three-channel color space, the combinations of these modalities in HiDiver has the potential to define a great number of new synthetic computational colors and stains.

#### Tractography and MRH resolution in relation to throughput

The long scan times required to generate 15 μm MRH data are excessive—currently a minimum of 5 days (**Table 1**). This prevents the acquisition of hundreds of scans per year. The primary ways to further reduce scan times are to: **1.** decrease spatial resolution, **2.** decrease angular resolution, **3.** increase the compression factor, or **4.** decrease the repetition rate. We chose to fix the *b* values—that is the magnitude of diffusion weighting—for all acquisitions at 3000 s/mm2 based on previous experiments (*Methods*)[32]. This imposes a limit on TR-dependence on the thermal load to the gradients. We have previously explored improving throughput by increasing compression factors [11], and while there may still be room for improvements, in our view the impact of further compression and acceleration is limited. The most practical route for continued reductions in scan times is optimization of spatial and angular resolution. Numerous papers have explored the tradeoffs between angular and spatial resolution, scan time, and signal-to-noise ratios, but not at this scale. Experiment 2 was designed to compare tradeoffs between the spatial and angular resolution with comparable scan times. We summarize the tradeoffs in **Methods**. **Figure S3** includes images of DWI and QA using a more efficient protocol (42 h) that is now being used in imaging genetic studies of aging and Alzheimer’s disease in the BXD and AD-BXD families [33].

#### LSM alignment to super-resolution MRH tract density images

We computed whole brain tractography in DSI Studio by seeding 4D MRH volumes with random sets of ~54 million points [34]. Track density images were generated using the super resolution method developed by Calamante and colleagues [35, 36]. The method imposes a grid on the volume which can be smaller than the acquired voxels. It takes advantage of the underlying model that generates the tracks by generating the density of tracks in these smaller grid elements derived from the whole brain tracking. The high contrast yellow strands in these TDIs are track densities binned with a super-resolution of 5 μm (see *Methods*). A crucial element in HiDiver is registering the TDI with the spatially corrected LSM images. The precision of registration across these radically different imaging modalities is striking (**Fig 4**). Despite the non-uniform distortions highlighted in **Fig 3**, we are able to achieve alignment that is usually at the level of cytoarchitectonic boundaries and cell layers. In many cases the alignment is almost at the level of single cells (**Fig 4C** and **F)**.

**Figure 4.**
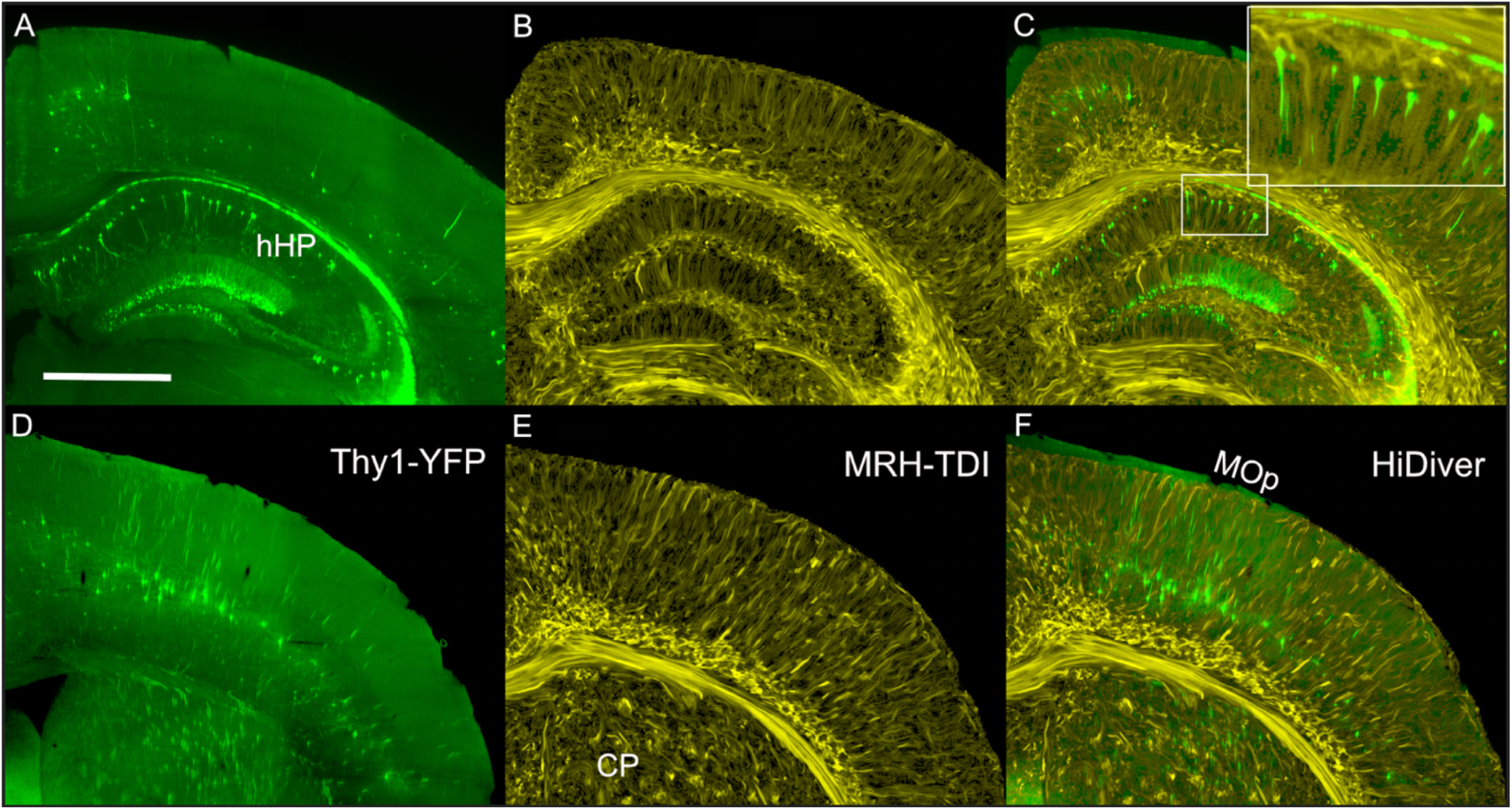
Joint LSM and super-resolution TDI at two levels through cortex, dorsal hippocampus, and caudoputamen. **A** and **D** are Thy1 fluorescence images (green) from a 90-day-old B6.Cg-Tg(Thy1-YFG)HJrs/J (sample 190415-2:1). **B** and **E.** TDI at the same levels at a super-resolution of 5 μm. **C** and **F** are merged HiDiver images that highlight the alignment of TDI and Thy1-positive pyramidal cells, dentate gyrus granule cells, and axon fascicles penetrating the caudoputamen. Inset in **C** is a 3X magnification of a radial section of CA1. **D, E,** and **F** are LSM, TDI, and HiDiver at the level of the primary motor cortex (MOp). All images have been rendered at the same effective slice thickness of 14.4 μm. Scale bar in **A** is 1 mm.

The advantage of joint LSM and super-resolution TDI is that we are better able to interpret the fine-structure resolved in **Figs 4B** and **E** compared to previous state-of-the-art work (e.g., figure 5C from [36]) In corpus callosum and other tracts. The fine strands are consistent with fascicles of myelinated axons that contrast with darker interfascicular callosal oligodendrocytes [37] [38] (**Fig 4B** and **E**). Some of the strands are nearly 1 mm long and have an apparent diameter of 15–30 μm. This is an order of magnitude more than the diameter of single large myelinated fiber—about 3 μm [39]. Another striking feature of the strands is their radial uniformity in fortuitous planes through cortex (**Fig 4E**). As strands reach the superficial cortical layers they splay out to form arcades, just as expected of distal dendrites of pyramidal cells and ascending axonal projections and collaterals (**Fig 4B, Fig S4**). Similarly, the fibrillar columns and arcades traversing hippocampal layers are probably heterogeneous but adjacent bundles of apical dendrites, axons, and glial processes (**Fig 4C,F**).

**Figure 5.**
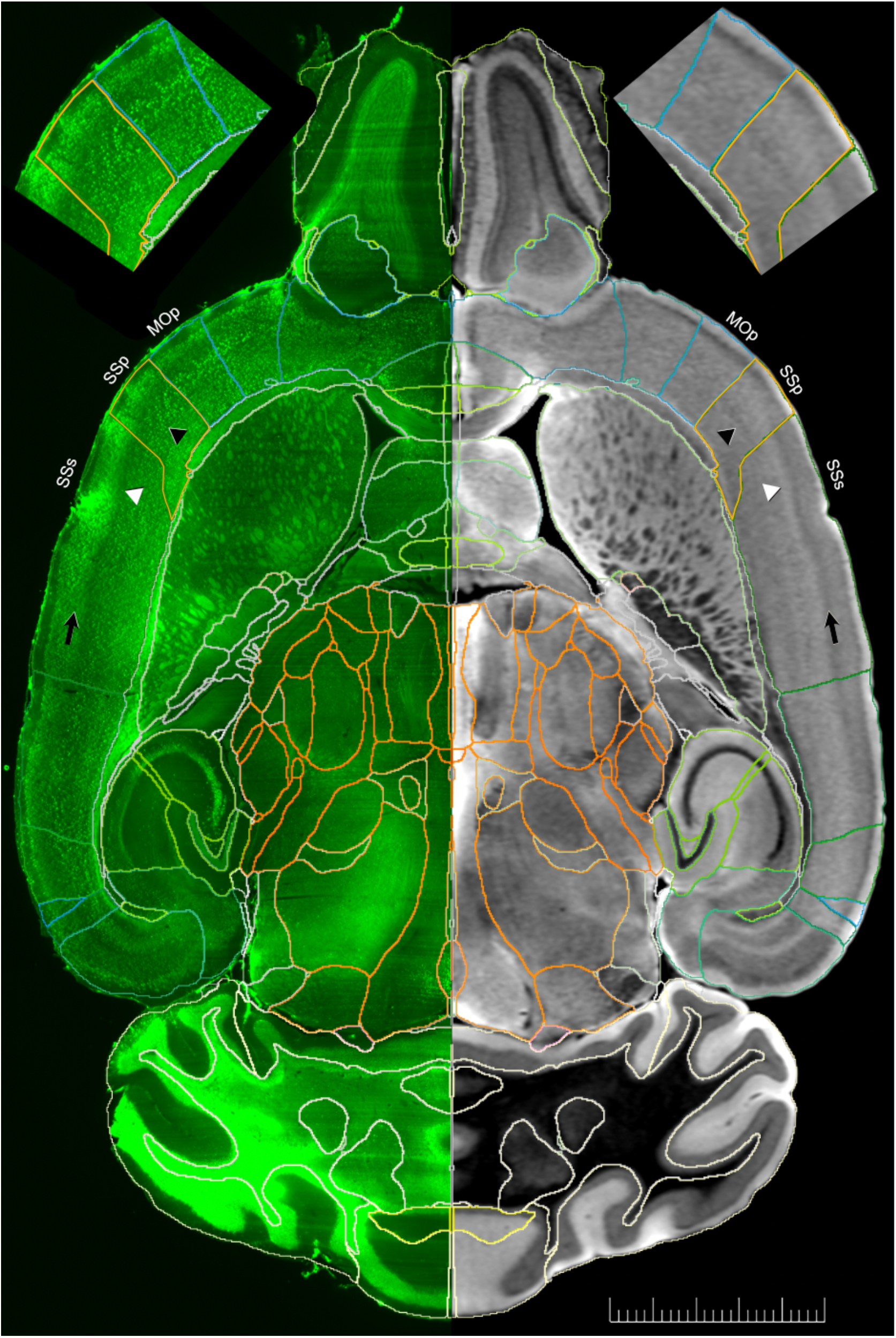
Accurate transfer of ROIs across age and genotype. NeuN (**left**) and axial DWI (**right**) and images are from an old BXD89 male (687 days, specimen 00803-12:1, also see **Fig 6B** and **D**). ROIs were lifted over from the C57BL/6J reference volume to the MRH using the SAMBA pipeline. ROIs were then transferred to the NeuN LSM channel using a second registration pipeline also built on ANTs. The final ROIs have been integrated into all three full resolution LSM (1.8 × 1.8 × 4.0 μm) volumes, such as the NeuN channel shown left. Three cortical areas are marked—the primary motor area (**MOp**), the primary somatosensory area (**SSp**), and the supplemental (secondary) somatosenory area (**SSs**). The **insets** above are 1.5X magnifications of these three areas. While LSM has higher resolution, the MRH DWI modality highlights cytoarchitectonic boundaries more prominently. The dark DWI band (**arrows**) in SSs is layer 4 whereas the adjacent light band is layer 5a (**white triangles**). In SSp, layer 4 is broadened whereas the deeper layer 5a is accentuated. In comparison, layer 2 is accentuated in MOp (**black triangles**). Here the comparison across MRH and LSM is particularly informative. For details on structure and function of these regions see [43] for MOp and [44] for SSp. **Scale bar** is 2.5 mm.

**Figure 6.**
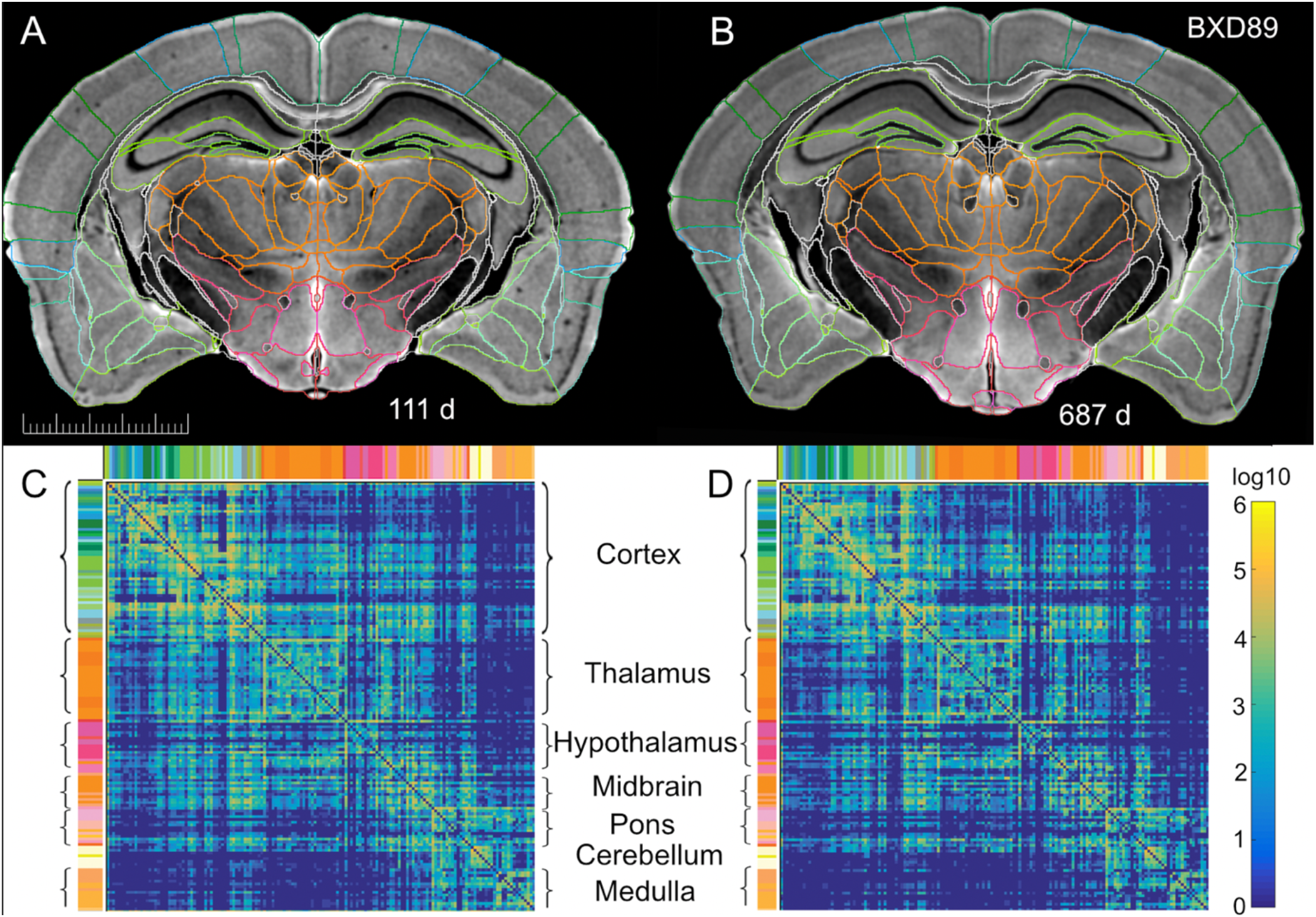
Accuracy of r1CCFv3 and quantitative tractography applied to pair of young and old BXD89 males. **A** and **B** are matched coronal images of cases that differ by 19 months and three generations). The alignment is more than satisfactory for quantitative analyses and comparison across genotype and age. **C** and **D** are matched quantitative connectomes of the same young and old cases that provide a metric (log10 scale) of the DTI connection strength of each of 152 selected right hemisphere gray matter ROIs with all other right hemisphere gray matter ROIs. The similarity of these matrices and the goodness of fit shown in A and B of ROI boundaries demonstrates the ability of HiDiver to follow subtle changes with age at a regional level.

To estimate effective resolution and alignment of HiDiver, we used the Thy1-labeled CA1 pyramidal cell bodies and their apical dendrites as a rough metric (**inset** in **Fig 4C)**. The diameters of cell bodies in the inset range from 20–30 μm, and apical dendrites taper from 8 to 4 μm. Individual TDI pixels in the inset that are 5 μm on a side can just be resolved in the inset— consistent with diameters of dendrites.

In summary, TDI provides excellent global metric of neuritic architecture of the CNS, and while it does not, in itself, distinguish among fascicle of axons, apposed dendrites, glial processes, or perhaps even small diameter vascular beds, this modality nonetheless represents a transformative new perspective on brain structure. In fact, these super-resolution 3D TDI data sets are essentially empirical full-volume versions of Krieg’s outstanding illustrations of the human brain (his figures 376–401 in [40]) but now made real at any plane or curvature of section in any mouse of any age.

#### Embedding full 3D volumetric labels from MRH into LSM

The next step in **experiment 2** was to lift over the entire set of volumetric delineations—bilaterally and independently—to the corrected LSM volume. This step is critical to enable global and regional cytometry of many classes of cells and neurons in hundreds of ROIs—one of the main goals of neuroscience [41], and the focus of our own neurogenetic dissection of the genetic control of regional size and cell numbers in CNS [42] [31]. But this step is also a computationally demanding task. The HiDiver lift-over of labels from MRH to LSM is good but still imperfect. The correspondence of the mirror-imaged horizontal planes**—**DWI (**Fig 5**, **left**) and NeuN (**right**)**—**is sufficiently accurate as judged by overall concordance of independently defined boundaries, blood vessels, and cytoarchitectonic boundaries. Interactive inspection enables panning and zooming through all the volumes. Vessels provide useful landmarks for comparison. In a substantial number of regions, the overlap is < 3voxels. In the case of the LSM and IHC staining, we note that penetration can occasionally be a challenge, but this is more of an issue with respect to cytometry than volumetrics and tractography. What we can assert now with confidence is that HiDiver—even at its first iteration—is sufficiently accurate to enable global quantification, and even cytometry of well over 180 ROIs. The technical caveat is that reagents must be able to penetrate efficiently 3–4 mm deep into brain tissue. Intrinsic transgenes should be less problematic, but these resources are somewhat sensitive to genetic background effects and subtle differences in promoter efficiency.

### Experiment 3. Robust Alignment: Impact of Age and Strain Variation on HiDiver

Having established that HiDiver can successfully register LSM with MRH and with r1CCFv3 segmentation, we asked whether this workflow can accommodate a markedly different genotype of mouse and at both young and old ages. Note that all cases used in experiment 1 and 2 are fully inbred C57BL/6J mice, the only subtle genetic difference being the insertion of a transgene in experiment 2. To test the robustness of HiDiver across genotype and age, we lifted over the r1CCFv ROIs into two BXD89 mice that differ from C57BL/6J at about 1 in 1000 base pairs of DNA—the level of genetic variation typically between two humans. The r1CCFv3 labels were transferred from the male reference (specimen 200302-1:1) to the new strains using our dedicated pipeline for rodents [45] (see **Fig S5** and **Methods** section on **Computation**). This code takes advantage of multiple contrasts generated by the DTI by using cross-correlation with DWI for the affine alignment and FA for the diffeomorphic registration. The transforms are then inverted to map ROIs from the reference to new cases. Transformations are applied to the labels, but crucially not to the underlying image data, thereby avoiding the potential of corrupting data. The HIDiver transfer of volumetric labels from r1CCFv3 to both BXD89 cases was remarkably accurate as judged by the boundaries imposed on the axial DWI MRH modality and on the corresponding LSF channels (**Figs 6,5**). To our eyes, the accuracy of boundaries is on par with that of the source C57BL/6J reference space and is generally well within 50 μm of the boundary as gauged by direct inspection of MRH and LSM images.

**We generated** whole brain connectomes by seeding 5 million points distributed across all ROIs (**Fig 6D, E**). For simplicity, only the connectomes of the right hemisphere are shown for young and old animals. These connectomes promise to be one of the most revealing phenotypes derived from HiDiver MRH workflow [31]. Maier-Hein and colleagues have highlighted the technical problems in human connectivity studies and the complications of false positive connections [46]. This is caused in large part by the modest spatial resolution of human connectomes—voxels of 1.25 to 2 mm3 [47]. These voxels can include well over overlapping 10,000 cells and axonal processes. In contrast, connectomes in **figure 6C** and **D** have resolutions 0.5 to 2.5 million times higher with commensurate reduction in accurately deconvolving the directions of fascicles [15]. The advantage of isogenic replication allows one to use powerful statistical tools to further refine the quality of data derived from these connectomes [31].

## DISCUSSION

### Synopsis of HiDiver

In summarizing the first century of neuroscience, Floyd E. Bloom credited Stanley Yolles with a mutated line from GB Shaw’s *Pygmalion*—“the gains in brain are mainly in the stain”. [48] Half a century later this still resonates, but now many of our “stains” are for genes, molecules, synapses, single cells, circuits, and connectomes. Progress on the fine-structural front is exemplified by innumerable technical and conceptual innovations in IHC, ISH, electron and light microscopic innovations, and single-molecular and single-cell labeling techniques.

The extensions and merging of staining and histological methods described here rely on a fundamental redesign of hardware and software infrastructure that now enables the collection and joint registration of key structural data for complete individual brains. This type of linkage is required to efficiently probe the gene-structure-function relations far much more systematically than has been possible to date. With HiDiver we can tie together cellular and subcellular structure with MRH modalities and with DTI connectomes of unprecedented resolution and geometric accuracy. A coherent quantitative analysis of regional volumes, cell and synapse types [49, 50], and single cell connectomes [51] is now practical as a global enterprise across thousands of unique genomes. For any one brain, the result is a comprehensive multimodal data package at a registration accuracy that is very close to the single cell level. And this package can be registered with behavioral measure for the same case. We believe this approach will offer exciting new ways to understand genetic variation and environmental perturbations that underpin differences in cells, circuits, and behaviors as a function of stage, age, and treatment.

### Significance of HiDiver

Each image modality has its own strengths and weakness in terms of accurate delimitation of regional boundaries and many other facets. The MRH modalities have excellent built-in multimodal registration, and high geometric fidelity to the *in vivo* brain. In other words, the MRH reference space is very close to a veridical brain volume. As a result, MRH defines histological features without appreciable distortion, shrinkage, or the registration challenges that compromises many histological procedures and that make stereological analyses challenging [25]. For example, classic celloidin and parafin embedding methods typically result in shrinkage of 50% or more [52, 53], whereas LSM can result in expansion of 50% or more. But in their defense light and electron microscopic methods obviously have orders of magnitude higher spatial resolution. What HiDiver now does is enable neuroscientists to efficiently and accurately pack multiple complementary data modalities into one combined virtual multimodal image that can be shared FAIRly [54]. The user can swim within a polymodal image pool, selecting viewports and magnifications at will. The exploration and analysis are limited only by the sophistication of the user interface, the speed of computing this virtual reality, and the imagination of the driver. HiDiver provides a framework for this work.

Our methods differ from previous work [55] [56] in five specific ways: **1.** the spatial resolution of MRH is improved by approximately three orders of magnitude; **2.** the contrast resolution is also higher, and there are multiple scalar images from the DTI/GQI acquisition that provide varied anatomical perspectives; **3.** MRH data are acquired in the skull with almost no distortion enabling LSM to be spatially remediated with excellent geometric accuracy with respect to *in vivo* morphometry; **4.** we provide a novel computational workflow that generates coherent multimodal packages of quantitative data for each specimen within a common and fully rendered reference space; and finally, **5**. the throughput is sufficiently high that all methods can be targeted at neurogenetic and genome-wide mapping studies which require large numbers of specimens [13, 57].

### Throughput and Challenges of HiDiver

HiDiver already has good throughput—sufficient for our group to have acquired over 100 MRH data sets at 45 μm resolution for over 20 strains during two intense years of methods development. **Figure S3** shows results from a recently streamlined protocol in which we have increased the spatial resolution from 45 μm to 25 μm while maintaining a throughput of three specimens per week. Current efforts are focused on a new strategy to accelerate acquisition by novel compression in Fourier (k) and diffusion (q) raw data space using modified key-hole sampling [58]. Use of magnetization preparation promises additional acceleration. We expect to be able to acquire 25 μm/61 angle data set in less than one day within the next year, a throughput that will support acquisition of hundreds of HiDiver data sets per year. LSM poses similar problem in terms of throughput and data volumes when using higher power objectives—20X and above and pixels of 0.2–0.5 μm.

### HiDiver and Genetic and Environmental Dissection of the Murine CNS

The conundrum now is how to snap together all these very large volumes of data in ways that can help answers the big questions related to structure-function mapping. Mere data integration, however elegant, is not a sufficient solution—it is just a key first step. What is also needed is a computational framework to systematically extract or test the causality of putative structure-function relations. As is typical in our reductionist age, almost all of neuroscience involving *C. elegans* and rodents still relies on single genotypes. The archetypal *C. elegans* is the N2 Bristol isogenic strain [59].The archetypal mouse is a 56-day-old C57BL/6J male [60], and the archetypal rat is a young Sprague Dawely male. This *n-of-1* design is neither computationally robust nor viable as a long-term solution with high translational relevance. It is not possible to use a single genome to derive structure-function relations that have any prospects of generalizing to hundreds of other members of the *Mus* and *Rattus* genuses, let alone to other members of the euarchontoglires superfamily of mammals that include rodents, all monkeys and apes, and of course humans.

### Near-term Prospects of MRH, HiDiver, and AI-driven Refinements

MRH modalities are inherently registered, and the entire processing workflow is remarkably repeatable with few artefacts. Consequently, MRH data sets provide a superb veridical framework for all methods of brain analysis. For example, in **Fig 1E**, the FA image is constructed from a normalized ratio of the AD and RD images resulting in an image that highlights white matter. Figure 1F takes that a step further by adding directional information derived from the eigen vectors of the voxel using red to delineate voxels in which the primary axis of diffusion within the voxel is along the left-right axis, green to encode diffusion along the anterior-posterior axis, and blue to encode diffusion along the superior-inferior axis. The result is easily appreciated in the three-color encoded image. But many more axes of information are available. The DTI algorithm yields a 3×3 tensor for each voxel from which the multiple scalars are derived. The GQI algorithm yields a much more comprehensive decomposition of the voxel into an orientation diffusion function from which multiple fibers can be isolated, each with magnitude and direction. This is exactly the sort of rich feature set on which AI algorithms flourish. It’s only a matter of time before many of the masking and registration steps we have used will be replaced by streamlined AI approaches. And these AI-driven workflow will yield hyperspectral cytoarchitectural vectors for each voxel covering the entire brain. What this also means is HiDiver will make it possible to bring LSM data in register with MRH within a cell diameter across the entire CNS. It should then become practical to construct vectors for each HiDiver voxel from multiple disparate modalities. These hyperspectral data could stream into algorithms to model the complex and recursive causal relations between genetic differences, age and sex differences complex structure and function, and multiple environmental factors and exposures.

As we further improve throughput, we anticipate that experimental studies will shift from a reductionist approach to more realistic and robust population models that incorporate levels of genetic variation comparable to human populations. With the power of the methods, we describe here, translation from murine models to humans will be both more mechanistically informative and clinically relevant.

## METHODS

The improvements in resolution, contrast, and throughput we report here are the culmination of nearly four decades of steady incremental progress. Paul Lauterbur closed his seminal paper on MRI with the statement that “zeugmatographic (imaging) techniques should find many useful applications in studies of the internal structures, states and composition of microscopic objects” [1] Magnetic resonance microscopy was subsequently demonstrated in 1986 by three groups [61] [62] [63] The application of MRI to the study of cytoarchitecture in fixed tissue, i.e. magnetic resonance histology (MRH) was suggested in 1993. MRH, while based on the same physical principals as MRI is fundamentally different than the clinical exams which are typically limited to v oxel dimensions of ~ 1 mm^3^. Preclinical imaging systems can acquire in vivo images with voxels ~ 1000 times smaller.

MRH allows us to extend the spatial resolution yet another factor of 1000. **Figure S6** is a comparison of a state-of-the-art FA images of a C57BL/6J brain *in vivo* at 150 μm resolution (voxel volume of 3.3 x10^−3^ mm^3^) with the *ex vivo* atlas we have generated for this work at 15 μm spatial resolution (voxel volume of 3.3 x 10^−6^ mm^3^). In previous work, we demonstrated the utility of MRH in neurogenetics at spatial/angular resolution of 45 μm with 46 angles [31]. At this spatial and angular resolution, it is possible to map connectomes with high correspondence to retroviral tracer studies [15]. But the MRH-derived connectomes can be generated in less than a day whereas global analysis with retroviral tracer can take years [64].

### Workflow

**Figure S5** provides an overview of the workflow—the tissue preparation, image acquisition pipelines for MRH, DTI, and LSM, the registration post-processing, and the final data integration into the common full volume rendered r1CCFv3 space. There are two main processing streams. The first focuses on the acquisition of 12 different MRH modalities with the brain in the cranial vault. The second focuses on the acquisition of multiple channels of LSM full brain 3D images. The two streams are merged into a common space define by the ABA CCF3v3 [65] to allow generation of quantitative image derived phenotypes of cells and circuits covering the whole brain.

### Specimen Preparation

All procedures were approved by the Duke University institution animal care and use committee. Three groups of animals were obtained. Adult male and female C57BL/6J mice were purchased from Jackson Laboratory for experiment 1 (**Table 1**). Adult male and female B6.Cg-Tg (Thy1-YFG/HJrs/J) mice were purchased from Jackson laboratory to validate tractography. BXD89 cases were obtained from the University of Tennessee Health Science Center to demonstrate the application of the methods in a study of the genetics of aging. All animals were allowed to adjust to their new environment for one week or more. All animals were perfused with 10% Prohance (Gadoteridol) in buffered formalin [66] [9] Prohance is a chelated gadolinium compound (Gd, a transition metal with unpaired electrons) commonly used in clinical MRI as a contrast agent. It is used as an active stain in MRH to reduce the spin lattice relaxation time (T1) from 1800 to 100 ms [66]. The animal was anesthetized to a surgical plane with Nembutal. A 21-gauge needle connected to a peristaltic pump was inserted in the left ventricle. Blood was flushed using a 0.1% heparin saline solution, followed by perfusion with the Prohance/formalin mixture for ~ 6 min. The head was placed in buffered formalin for an additional 24 h. The mandible was removed and the brain, still in the cranium, was placed in an 0.5% Prohance/buffer solution.

### MR Histology

MRH images were acquired on a 9.4T/89 mm vertical bore magnet controlled by an Agilent Direct Drive console (Vnmrj 4.0). The gradient insert we used was constructed by Resonance Research Inc (Billerica, Ma). The high-performance gradient (BRG-88_14) provides peak gradients up to 2500 mT/m with rise times of 25000 T/m/s and ±5% linearity over a 30 mm diameter spherical volume. The high-capacity cooling system (600 W power dissipation) allowed us to use exceptionally strong gradients (1500 mT/m) with high duty cycle (TR = 100 ms). The perfusion fixed brain is placed in a 10 mm diameter tube with a 3D printed insert to stabilize it in the container. The tissue is surrounded by fomblin, a fluorinated lubricant that minimizes the susceptibility artifacts at the surface of the brain. The container is placed in a solenoid coil constructed from a single sheet of silver which yields high Q (>600), high homogeneity, and high sensitivity. Two imaging sequences were used; 1) a multigradient echo sequence (TR/TE = 100/4.4 ms) and a Stesjkal/Tanner spin echo sequence for diffusion tensor imaging (DTI) (TR/TE =100/12-19 ms) [67]. Both sequences employ phase encoding along the short axes of the specimen (x and y) with the readout gradient applied along the long axis of the specimen (z).

### Computer Resources

Standard MRH workflows and HiDiver in particular require high-performance computer systems (**Figure S7**) that can handle multiple 3D arrays as large as 1 TB. In our implementation, source image files are streamed to a 604-core 18-node cluster for nearly automated computation and joint alignment. We have also relied on two servers with 1.5 TB of memory that enable interactive analysis of HiDiver data using software such as Fiji, Slicer, and Imaris. Large (100 TB) high performance RAIDs provide local storage to facilitate online interactive analysis via Citrix.

### Compressed Sensing Reconstruction

Compressed sensing is employed to accelerate both sequences [68] [11]. A probabilistic map was generated for sparse sampling of Fourier space along the two-phase encoding axes **(Fig S8)**. A script on the scanner starts the acquisition of the first 3D volume, a baseline image (b0) with no diffusion encoding gradient. At the conclusion of the scan, the data is automatically written to the cluster. A Fourier transform is applied along the z axis (the long axis of the brain) producing between 256–3000 individual 2D files which are distributed to multiple processors for parallel iterative reconstruction. When all the 2D files have been reconstructed, the data is reassembled into a 3D volume. While this is under way, the scanner launches the next volume in the sequence with a diffusion encoding direction (bvec) and value (bval) determined from a file accessed via the scanner script. The process proceeds under control of the scanner script through as many bvecs and bvals as one chooses. A baseline acquisition is included in every 10-15th volume to monitor any drift in the spectrometer.

### 4D volume creation

The strong gradients used for diffusion encoding induce eddy currents in the magnet bore that spatially shift the individual 3D volumes. The shifts are dependent on the magnitude and direction of the diffusion gradients. A skull stripping algorithm is applied to each volume as a preprocessing step that facilitates registration of the multiple 3D volumes into a single 4D array. An average baseline image (Avgb_0_) is created by registering the baseline images generated at regular intervals through the course of the scan **(Table S1)**. The pipeline based on ANTs uses scaling and affine transforms to produce the average baseline image. The DWI volumes are then registered to Avgb0 producing a 4D array of the registered 3D volumes [29] [45].

### Tradeoff in spatial and angular resolution

The propagation of error (noise) through the multiple pipelines (compressed sensing, registration, DSI Studio) is unclear. Experiment 2 provides details on three data sets acquired to experimentally compare these tradeoffs (**Table 1)**. The cycle repetition time (TR), echo time (TE) and *b* values were fixed for all three scans with attention to the maximum thermal loading for our gradients. Figure S9 shows a comparison of the FA images. Images have been scaled to the same size for comparison. The first two scans to compare 15 μm (**Fig S9A**) vs 25 μm (**Fig S9B**) spatial resolution was performed with approximately the same total scan time, the slight variation coming from the desire to choose angular sampling strategies in which the b vectors were equally spaced on the unit sphere. As one would expect, the 15 μm data is noisier than the 25 μm data. The definition of cortical layers and layers in the hippocampus is better in the lower resolution, higher SNR (25 μm) data. The impact of angular sampling i.e. comparison of **figure S9B** (at 126 angles) with **figure S9C** (at 61 angles) is not as obvious yet it results in a reduction of scan time by ~1/2. Recent work has demonstrated an inflection point at ~61 angles when comparing connectome metrics vs angular sampling [69].

### Denoising

We have introduced a denoising step that renders the noise in the scalar images of the 25 μm /61 angle array at comparable quality to 25 μm/126 angles. St-Jean et al have developed a novel denoising algorithm that recognizes the fourth (angular) dimension of the data [70]. The volume is decomposed into 4D overlapping patches that sample both the spatial and angular resolution. A dictionary of atoms is learned on the patches and a sparse decomposition is generated by bounding the reconstruction error with the local noise variance. The method improves the visibility of structures while simultaneously reducing the number of spurious tracts. St-Jean and colleagues implemented the algorithm for high resolution clinical scans (matrix of 210×210×210) of ~ 20 MB/volume with 40 volumes (~800 MB). The atlases generated for this work were acquired with arrays as large 800×800×1600 with 108 angular samples plus 13 baseline images i.e. a 4D volume (~252 GB) that is more than 300 times larger. To accommodate the change of scale, the algorithm was implemented on the cluster (**Fig S7**) by breaking the volume into overlapping cubes which could be processed in parallel**. Figure S10** shows the diffusion weighted images from specimen 200316-1:1 before and after denoising (**Table 1**).

### Generation of MRH Modalities

The four echoes of the MGRE image are averaged together to generate an (AvgMGRE) image in which tissue contrast is driven by the transverse relaxation time and local field inhomogeneity (T2*-weighted). A MATLAB script averages all the diffusion weighted 3D volumes to produce the diffusion weighted image (DWI). The 4D denoised volume is passed to DSI Studio (http://dsi-studio.labsolver.org/) where a perl script executes an initial pass using the diffusion tensor algorithm [26] to generate five different 3D scalar volumes: 1. axial diffusivity (AD), 2. radial diffusivity (RD), 3. mean diffusivity (MD), 4. fractional anisotropy (FA), and 5. the color fractional anisotropy (clrFA) images. These six 3D scalar images have decidedly different contrasts and highlight different anatomical features, boundaries, and orientations of cellular components (**Fig 1**). The combination of MGRE and DTI scalar data drive the registration of the atlas and associated labels.

### Label registration

The refence volume and associated labels are registered to the strain under study using our previously published SAMBA pipeline [45]. An initial rigid transformation aligns the unknown to the RCCF atlas described above. In population studies with multiple specimens of a given strain, a minimum deformation template (MDT) is formed through iterative affine and diffeomorphic transforms using multiple combinations of the scalar images. A combination of DWI and FA has proven effective for the studies we have undertaken to date. Once an MDT is complete, all specimens in that cohort are reregistered to it. Registration between images of similar contrast (FA to FA) are executed using the cross-correlation similarity metric. Registration between images of differing contrast (FA to DWI) use mutual information. When the atlas and labels have been mapped to the specimen under study, the transform is inverted to map the labels back to the specimen under study leaving a 4D volume with labels for the new specimen from which scalar image phenotypes can be derived. A MATLAB script extracts volume, and mean and standard deviation of the scalar diffusion metrics (AD, RD,MD,FA) for all 180 (x2) regions of interest. The pipeline calculates a coefficient of variation between left and right hemispheres as a quality assurance check. The summary of regional scalar image derived phenotypes is written to a summary spread sheet with the meta data for the study [31].

### Tractography and connectome generation

The 4D volume with labels, b vectors and b values are passed to DSI studio to generate whole brain tractography, track density images, connectome and associated statistics using the GQI algorithm. The GQI algorithm exploits the higher angular sampling resolving multiple fibers (up to 4) in each voxel [27, 34]. Whole brain tractography is executed on CTX01 (**Fig S6**), a server with 1.5TB of memory, sufficient for the large .fib file generated in DSI Studio. Connectomes are generated using the labels generated in the label r egistration pipeline using quantitative anisotropy (QA), the metric in the GQI algorithm analogous to FA in the DTI algorithm.

Track density images are generated in DSI Studio using a super sampling algorithm described by Calamante et al [35]. The algorithm has been previously validated against conventional histopathology [36]. Images in these previous studies were acquired with 100 μm spatial resolution, 30 angles, i.e. a resolution index of ~3.0×104 [16]. Using super resolution, Calamante extended the spatial resolution for the TDI to 20 μm by seeding the whole brain to 4 million fibers and tracking with a step size of 0.1 mm. In comparison, the resolution index achieved here is 3.2×10^7^ about three orders of magnitude higher than previous work. Thus, for our work, whole brain tractography was acquired in DSI Studio by seeding the 4D volume the brain with a random set of 54 million points. Step size was set to 0.75 μm at a QA threshold of 0.1% of the peak QA histogram generating track density images with nominal sampling of 5 μm. The resulting large arrays (TDI ~34 GB and LS ~200 GB) were examined in registration with MRH in Imaris (Oxford Instruments).

### LSM acquisition

The brain was extracted from the skull and placed in normal saline and shipped to LifeCanvas Technologies in an airtight container to minimize the formation of bubbles in the brain during air shipping. The tissue was cleared using SHIELD [71] and stained using SWITCH [71]. The smart SPIM light sheet fluorescence microscope provides whole brain coverage (3650 μm field of view) at 1.8 × 1.8 × 4.0 μm with a 3.6X objective producing three stacks of registered 2D .tiff images—one stack for each excitation wavelength. Typical array size for a stack is 7600 × 10600 × 2250 or about 250 GB.

### LSM registration

The three collections were transferred via Globus to the sever configured for light sheet image analysis (**Figure S6**). The NeuN channel was registered to the DWI for the atlas creation and aging comparisons. The autofluorescence images were registered to the DWI for the tractography comparisons. In step 1, the LSM volume was down sampled to the resolution of the MRH volume (15 μm or 25 μm). The LSM were passed through an initial preconditioning step to address the substantial distortion arising from the tissue handling. This required manual placement of ~20 landmarks in the areas that suffered the greatest distortion (brain stem and olfactory bulb). A pipeline built on ANTs was constructed to execute the registration in three stages: 1) Affine transformation using mutual information; 2)B spline using cross correlation; 3) Diffeomorphic transform using mutual information. The resulting transform from the low resolution (15 or 25 μm) arrays were then applied to the full resolution LSM images.

## ACKNOWLEDGEMENTS

Our special thanks to Dr. Glenn D. Rosen for invaluable help in developing the r1CCFv3 used for the first time in this publication. Thanks to Drs. Lu Lu, Suheeta Roy, and David G. Ashbrook for access to BXD mice and aged animals from the UTHSC Aging Colony. Thanks to Peter Nicholls for the Golgi image in **Figure S4**. We are grateful to Lucy Upchurch for invaluable technical assistance. We thank Tatiana Johnson for special care in manuscript preparation and submission.

## SUPPLEMENTARY MATERIALS

Supplementary material associated with this article can be found in the online version at doi: ∗Corresponding author at: Dr. G. Allan Johnson, Duke Center for In Vivo Microscopy, Department of Radiology, Duke University, Duke University Medical Center Box 3302, Durham, NC 27710, USA. *E-mail addresses:*

## AUTHOR CONTRIBUTIONS

GAJ led the project, directed data acquisition, development of r1CCF3, analyses, figure preparation, and writing; GPC developed the MR pulse sequences (with NW) for compressed sensing and acquired the MR data; JCC- developed and maintains the imaging pipelines, assisted in data processing; JCG- assisted in translating CCF3 labels to MR atlas; AH assisted in acquisition of the LSM data provided direction in choice of immune histochemical labels; KH assisted in data processing and figure preparation; YQ managed brain tissue preparation and dissection for LSM; YT developed (with JCC) the LSM registration pipeline and assisted in data processing; FCY handled data sets in DSI Studio and assisted in adapting code for HiDiver processing; NW developed the compressed sensing pipeline and assisted in MR acquisition; LEW assisted in data interpretation, development of the r1CCFv3 atlas, neuroanatomical interpretation, and writing; RWW helped in conception, production of the r1CCF1 atlas with Dr. GD Rosen, HiDiver data integration and analyses, neuroanatomical interpretation, figure preparation, and writing.

