## Supplementary Figures and Supplemental Table 1 for "HiDiver: A Suite of Methods to Merge Magnetic Resonance Histology, Light Sheet Microscopy, and Complete Brain Delineations"

**Supplemental Figures**

| **Source Data** | **Algorithm** | **Name** | **Abbreviation** |
| --- | --- | --- | --- |
| MGRE | Average | Average | AvgMGRE |
| DTI | ANTs | Average Baseline | Avgb_0_ |
|  | Average | Diffusion Weighted Image | DWI |
|  | DTI | Mean Diffusivity | MD |
|  |  | Axial Diffusivity | AD |
|  |  | Radial Diffusivity | RB |
|  |  | Fractional Anisotropy | FA |
|  |  | Color Fractional Anisotropy | ClrFA |
| DTI | GQI | Isotropic Fraction | iso |
|  |  | Normalized Quantitative Anisotropy | nqa |
|  | Calamante | Track Density Imaging | TDI |
|  |  | Color Track Density Imaging | ClrTDI |

Table S1 Summary of imaging modalities generated in the standard HiDiver/MRH protocol

including the algorithms used and the abbreviations

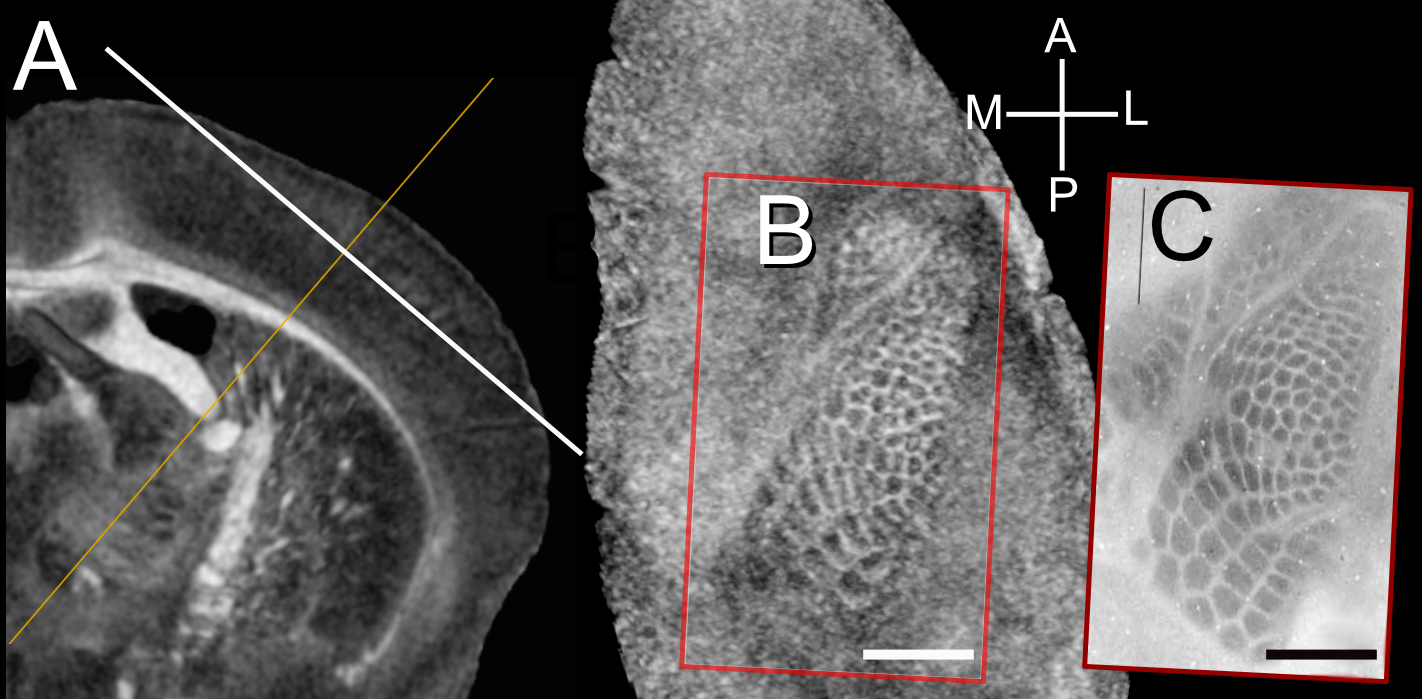

**Fig S1. *Barrel cortex digital flatmount of a C57BL/6J*** *(Specimen 200302-1:1).* ***A.*** *Coronal FA section in which the orientation of the tangential plane through layer 4 of the barrel field is highlighted by a long white two-headed arrow.* ***B.*** *A single 15-µm-thick DTI image slice of the left cortex in a plane tangentially to the barrel field. The white calibration bars is 1 mm. In* ***C*** *we have juxtaposed an image of a cortical flatmount from the same strain taken from [1] (their figure 1B with its own thin black 1 mm calibration bar). The alignment is remarkably precise with only linear rescaling.*

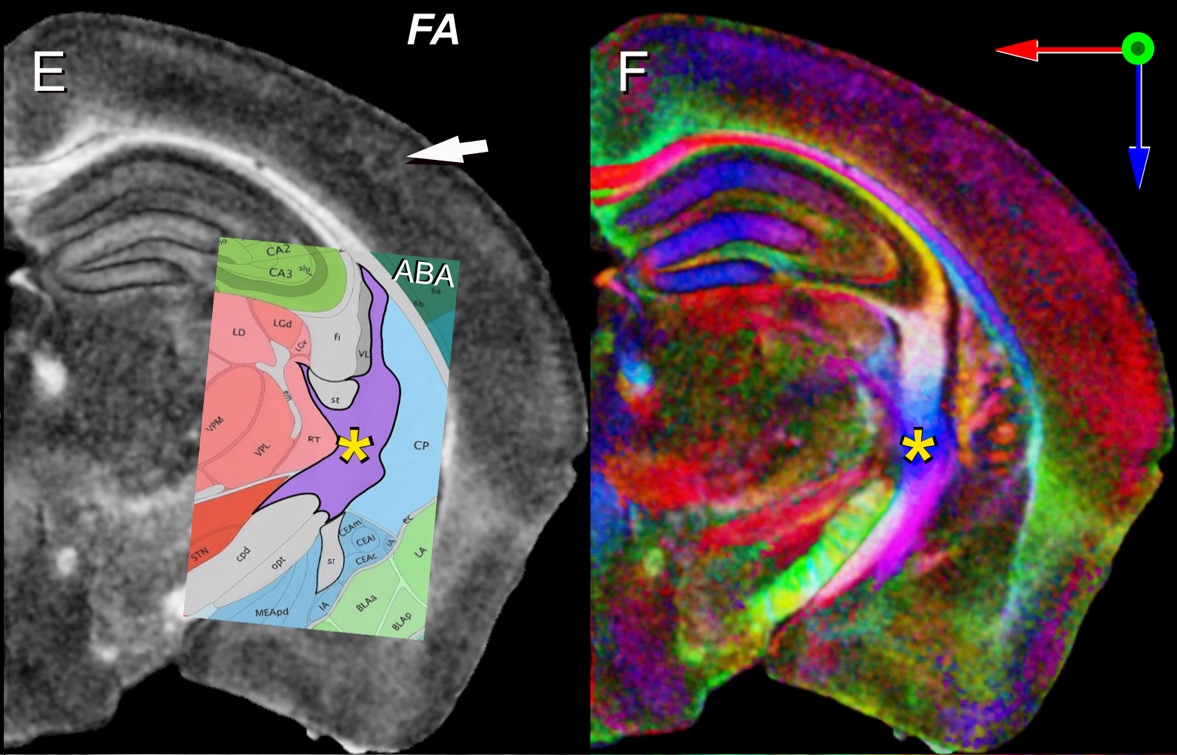

***Figure S2.*** *This is an extension of Figure 1.* ***Panel E*** *displays the fractional anisotropy where image intensity reflects the anisotropic diffusion in each voxel****.*** *The overlay in panel* ***E*** *is from the Allen Brain Atlas CCFv3 [2].****Panel F*** *adds color in which the primary axis of diffusion is encoded in color. It is straightforward in the color FA to define all tracts disambiguated in CCFv3.*

| **ROI#** | **Name** | **Acronym** | **CCF3 Structures Combined** |
| --- | --- | --- | --- |
| 1 | Cerebrum | CH |  |
| 2 | Basic cell groups and regions | grey |  |
| 3 | Olfactory bulb | OB | AOB, AON, MOB |
| 4 | Anterior olfactory nucleus | AON |  |
| 5 | Frontal pole, cerebral cortex | FRP |  |
| 6 | Orbital area | ORB |  |
| 7 | Prelimbic area | PL |  |
| 8 | Infralimbic area | ILA |  |
| 9 | Anterior cingulate area | ACA |  |
| 10 | Agranular insular area | AI |  |
| 11 | Gustatory areas | GU |  |
| 12 | Piriform area | PIR |  |
| 13 | Visceral area | VISC |  |
| 14 | Primary motor area | MOp |  |
| 15 | Secondary motor area | MOs |  |
| 16 | Primary somatosensory area | SSp |  |
| 17 | Primary somatosensory area, barrel field | SSp-bfd |  |
| 18 | Supplemental somatosensory area | SSs |  |
| 19 | Posterior parietal association areas | PTLp |  |
| 20 | Auditory areas | AUD |  |
| 21 | Temporal association areas | TEa |  |
| 22 | Ectorhinal area | ECT |  |
| 23 | Perirhinal area | PERI |  |
| 24 | Retrosplenial area | RSP |  |
| 25 | Visual areas | VIS |  |
| 26 | Primary visual area | VISp |  |
| 27 | Entorhinal area | ENT |  |
| 28 | Subicular region | SUBR | SUB, PRE, POST, PAR, ProS |
| 29 | Hippocampus | HI | CA, DG |
| 30 | Hippocampal region | HIP |  |
| 31 | Field CA1 | CA1 |  |
| 32 | Field CA3 | CA3 |  |
| 33 | Dentate gyrus | DG |  |
| 34 | Dentate gyrus, granule cell layer | DG-sg |  |
| 35 | Dentate gyrus, polymorph layer | DG-po |  |
| 36 | Claustrum | CLA |  |
| 37 | Central amygdalar nucleus | CEA |  |
| 38 | Striatum-like amygdalar nuclei | sAMY |  |
| 39 | Medial amygdalar nucleus | MEA |  |
| 40 | Lateral amygdalar nucleus | LA |  |
| 41 | Basolateral amygdalar nucleus | BLA |  |
| 42 | Basomedial amygdalar nucleus | BMA |  |
| 43 | Posterior amygdalar nucleus | PA |  |
| 44 | Cortical amygdalar zones | COAz | COA, PAA, TR |
| 45 | Endopiriform nucleus | EP |  |
| 46 | Striatum | STR |  |
| 47 | Striatum dorsal region | STRd |  |
| 48 | Nucleus accumbens | ACB |  |
| 49 | Fundus of striatum | FS |  |
| 50 | Olfactory tubercle | OT |  |
| 51 | Globus pallidus, external segment | GPe |  |
| 52 | Globus pallidus, internal segment | GPi |  |
| 53 | Substantia innominata | SI |  |
| 54 | Medial septal complex | MSC |  |
| 55 | Lateral septal nucleus | LS |  |
| 56 | Bed nuclei of the stria terminalis | BST |  |
| 57 | Magnocellular nucleus _Meynert_ | MA |  |
| 58 | Thalamus | TH |  |
| 59 | Anteromedial nucleus | AM |  |
| 60 | Anterodorsal nucleus | AD |  |
| 61 | Anteroventral nucleus of thalamus | AV |  |
| 62 | Lateral dorsal nucleus of the thalamus | LD |  |
| 63 | Ventral anterior-lateral complex of the thalamus | VAL |  |
| 64 | Ventral medial nucleus of the thalamus | VM |  |
| 65 | Ventral posterior complex of the thalamus | VP |  |
| 66 | Ventral posteromedial nucleus of the thalamus | VPM |  |
| 67 | Ventral posterolateral nucleus of the thalamus | VPL |  |
| 68 | Parataenial nucleus | PT |  |
| 69 | Paraventricular nucleus of the thalamus | PVT |  |
| 70 | Nucleus of reuniens | RE |  |
| 71 | Central lateral nucleus of the thalamus | CL |  |
| 72 | Central medial nucleus of the thalamus | CM |  |
| 73 | Paracentral nucleus | PCN |  |
| 74 | Parafascicular nucleus | PF |  |
| 75 | Posterior intralaminar thalamic nucleus | PIL |  |
| 76 | Mediodorsal nucleus of thalamus | MD |  |
| 77 | Lateral posterior nucleus of the thalamus | LP |  |
| 78 | Posterior complex of the thalamus | PO |  |
| 79 | Medial geniculate complex, dorsal part | MGd |  |
| 80 | Medial geniculate complex, ventral part | MGv |  |
| 81 | Medial geniculate complex, medial part | MGm |  |
| 82 | Dorsal part of the lateral geniculate complex | LGd |  |
| 83 | Ventral part of the lateral geniculate complex | LGv |  |
| 84 | Reticular nucleus of the thalamus | RT |  |
| 85 | Lateral habenula | LH |  |
| 86 | Medial habenula | MH |  |
| 87 | Hypothalamus | HY |  |
| 88 | Medial preoptic nucleus | MPN |  |
| 89 | Anterior hypothalamic nucleus | AHN |  |
| 90 | Ventromedial hypothalamic nucleus | VMH |  |
| 91 | Posterior hypothalamic nucleus | PH |  |
| 92 | Mammillary body | MBO |  |
| 93 | Suprachiasmatic nucleus | SCH |  |
| 94 | Arcuate hypothalamic nucleus | ARH |  |
| 95 | Paraventricular hypothalamic nucleus | PVH |  |
| 96 | Lateral preoptic area | LPO |  |
| 97 | Tuberal nucleus | TU |  |
| 98 | Lateral hypothalamic area | LHA |  |
| 99 | Subthalamic nucleus | STN |  |
| 100 | Zona incerta | ZI |  |
| 101 | Superior colliculus, sensory related | SCs |  |
| 102 | Superior colliculus, motor related | SCm |  |
| 103 | Inferior colliculus | IC |  |
| 104 | Inferior colliculus, central nucleus | ICc |  |
| 105 | Parabigeminal nucleus | PBG |  |
| 106 | Pedunculopontine nucleus | PPN |  |
| 107 | Pretectal region | PRT |  |
| 108 | Oculomotor region | IIIR | III, EW |
| 109 | Periaqueductal gray | PAG |  |
| 110 | Red nucleus | RN |  |
| 111 | Midbrain reticular nucleus | MRN |  |
| 112 | Cuneiform nucleus | CUN |  |
| 113 | Interpeduncular nucleus | IPN |  |
| 114 | Ventral tegmental area | VTA |  |
| 115 | Substantia nigra, compact part | SNc |  |
| 116 | Substantia nigra, reticular part | SNr |  |
| 117 | Dorsal nucleus raphe | DR |  |
| 118 | Pons | P |  |
| 119 | Nucleus of the lateral lemniscus | NLL |  |
| 120 | Superior olivary complex | SOC |  |
| 121 | Parabrachial nucleus | PB |  |
| 122 | Principal sensory nucleus of the trigeminal | PSV |  |
| 123 | Pontine central gray | PCG |  |
| 124 | Pontine gray | PG |  |
| 125 | Trigeminal motor complex | VMc | V, P5, I5, Acs5 |
| 126 | Facial motor nucleus | VII |  |
| 127 | Dorsal tegmental nucleus | DTN |  |
| 128 | Laterodorsal tegmental nucleus | LDT |  |
| 129 | Tegmental reticular nucleus | TRN |  |
| 130 | Pontine reticular formation | PRF | PRNc, PRNr |
| 131 | Raphe ventral complex | RVC | IF, IPN, RL, CLI, CS |
| 132 | Hemispheric regions | HEM |  |
| 133 | Vermal regions | VERM |  |
| 134 | Paraflocculus | PFL |  |
| 135 | Floccular nodular region | FLNOD | NOD, FL, UVU |
| 136 | Dentate nucleus | DN |  |
| 137 | Interposed nucleus | IP |  |
| 138 | Fastigial nucleus | FN |  |
| 139 | Vestibulocerebellar nucleus | VeCB |  |
| 140 | Nucleus of the trapezoid body | NTB |  |
| 141 | Spinal nucleus of the trigeminal region | SPVr | SPVC, SPVI, SPVO |
| 142 | Nucleus of the solitary tract | NTS |  |
| 143 | Dorsal cochlear nucleus | DCO |  |
| 144 | Ventral cochlear nucleus | VCO |  |
| 145 | Vestibular nuclei | VNC |  |
| 146 | Spinal vestibular nucleus | SPIV |  |
| 147 | Dorsal column nuclei | DCN |  |
| 148 | Inferior olivary complex | IO |  |
| 149 | Perihypoglossal nuclei | PHY |  |
| 150 | Dorsal motor nucleus of the vagus nerve | DMX |  |
| 151 | Hypoglossal nucleus | XII |  |
| 152 | Medullary reticular zone | MRz | GRN, LRN, MARN, MDRN, PARN, PGRN, |
| 153 | Intermediate reticular nucleus | IRN |  |
| 154 | Medulla, raphe, behavioral state related | MY-sat |  |
| 155 | fiber tracts | fiber tracts |  |
| 156 | corpus callosum | cc | ccb, ccg, ccs, fp |
| 157 | cingulum bundle | cing |  |
| 158 | anterior commissure | ac | act, aco |
| 159 | optic tract and chiasm | optc | opt,och |
| 160 | dorsal hippocampal commissure | dhc |  |
| 161 | fimbria | fi |  |
| 162 | columns of the fornix | fx |  |
| 163 | stria terminalis | st |  |
| 164 | thalamus related | lfbst |  |
| 165 | medial forebrain bundle | mfb |  |
| 166 | fasciculus retroflexus | fr |  |
| 167 | posterior commissure | pc |  |
| 168 | corticospinal tract | cst |  |
| 169 | medial lemniscus | ml |  |
| 170 | lateral lemniscus | ll |  |
| 171 | medial longitudinal fascicle | mlf |  |
| 172 | superior cerebelar peduncles | scp |  |
| 173 | inferior cerebellar peduncle | icp |  |
| 174 | trigeminal nerve | Vn |  |
| 175 | cochlear nerve | cVIIIn |  |
| 176 | lateral ventricle | VL |  |
| 177 | third ventricle | V3 |  |
| 178 | fourth ventricle | V4 |  |
| 179 | cerebral aqueduct | AQ |  |
| 180 | Whole brain _root_ | Brain |  |
| 1001 | Cerebrum | CH |  |
| 1002 | Basic cell groups and regions | grey |  |
| 1003 | Olfactory bulb | OB | AOB, AON, MOB |
| 1004 | Anterior olfactory nucleus | AON |  |
| 1005 | Frontal pole, cerebral cortex | FRP |  |
| 1006 | Orbital area | ORB |  |
| 1007 | Prelimbic area | PL |  |
| 1008 | Infralimbic area | ILA |  |
| 1009 | Anterior cingulate area | ACA |  |
| 1010 | Agranular insular area | AI |  |
| 1011 | Gustatory areas | GU |  |
| 1012 | Piriform area | PIR |  |
| 1013 | Visceral area | VISC |  |
| 1014 | Primary motor area | MOp |  |
| 1015 | Secondary motor area | MOs |  |
| 1016 | Primary somatosensory area | SSp |  |
| 1017 | Primary somatosensory area, barrel field | SSp-bfd |  |
| 1018 | Supplemental somatosensory area | SSs |  |
| 1019 | Posterior parietal association areas | PTLp |  |
| 1020 | Auditory areas | AUD |  |
| 1021 | Temporal association areas | TEa |  |
| 1022 | Ectorhinal area | ECT |  |
| 1023 | Perirhinal area | PERI |  |
| 1024 | Retrosplenial area | RSP |  |
| 1025 | Visual areas | VIS |  |
| 1026 | Primary visual area | VISp |  |
| 1027 | Entorhinal area | ENT |  |
| 1028 | Subicular region | SUBR | SUB, PRE, POST, PAR, ProS |
| 1029 | Hippocampus | HI | CA, DG |
| 1030 | Hippocampal region | CA3 |  |
| 1031 | Field CA1 | CA1 |  |
| 1032 | Field CA3 | CA3 |  |
| 1033 | Dentate gyrus | DG |  |
| 1034 | Dentate gyrus, granule cell layer | DG-sg |  |
| 1035 | Dentate gyrus, polymorph layer | DG-po |  |
| 1036 | Claustrum | CLA |  |
| 1037 | Central amygdalar nucleus | CEA |  |
| 1038 | Striatum-like amygdalar nuclei | sAMY |  |
| 1039 | Medial amygdalar nucleus | MEA |  |
| 1040 | Lateral amygdalar nucleus | LA |  |
| 1041 | Basolateral amygdalar nucleus | BLA |  |
| 1042 | Basomedial amygdalar nucleus | BMA |  |
| 1043 | Posterior amygdalar nucleus | PA |  |
| 1044 | Cortical amygdalar zones | COAz | COA, PAA, TR |
| 1045 | Endopiriform nucleus | EP |  |
| 1046 | Striatum | STR |  |
| 1047 | Striatum dorsal region | STRd |  |
| 1048 | Nucleus accumbens | ACB |  |
| 1049 | Fundus of striatum | FS |  |
| 1050 | Olfactory tubercle | OT |  |
| 1051 | Globus pallidus, external segment | GPe |  |
| 1052 | Globus pallidus, internal segment | GPi |  |
| 1053 | Substantia innominata | SI |  |
| 1054 | Medial septal complex | MSC |  |
| 1055 | Lateral septal nucleus | LS |  |
| 1056 | Bed nuclei of the stria terminalis | BST |  |
| 1057 | Magnocellular nucleus _Meynert_ | MA |  |
| 1058 | Thalamus | TH |  |
| 1059 | Anteromedial nucleus | AM |  |
| 1060 | Anterodorsal nucleus | AD |  |
| 1061 | Anteroventral nucleus of thalamus | AV |  |
| 1062 | Lateral dorsal nucleus of the thalamus | LD |  |
| 1063 | Ventral anterior-lateral complex of the thalamus | VAL |  |
| 1064 | Ventral medial nucleus of the thalamus | VM |  |
| 1065 | Ventral posterior complex of the thalamus | VP |  |
| 1066 | Ventral posteromedial nucleus of the thalamus | VPM |  |
| 1067 | Ventral posterolateral nucleus of the thalamus | VPL |  |
| 1068 | Parataenial nucleus | PT |  |
| 1069 | Paraventricular nucleus of the thalamus | PVT |  |
| 1070 | Nucleus of reuniens | RE |  |
| 1071 | Central lateral nucleus of the thalamus | CL |  |
| 1072 | Central medial nucleus of the thalamus | CM |  |
| 1073 | Paracentral nucleus | PCN |  |
| 1074 | Parafascicular nucleus | PF |  |
| 1075 | Posterior intralaminar thalamic nucleus | PIL |  |
| 1076 | Mediodorsal nucleus of thalamus | MD |  |
| 1077 | Lateral posterior nucleus of the thalamus | LP |  |
| 1078 | Posterior complex of the thalamus | PO |  |
| 1079 | Medial geniculate complex, dorsal part | MGd |  |
| 1080 | Medial geniculate complex, ventral part | MGv |  |
| 1081 | Medial geniculate complex, medial part | MGm |  |
| 1082 | Dorsal part of the lateral geniculate complex | LGd |  |
| 1083 | Ventral part of the lateral geniculate complex | LGv |  |
| 1084 | Reticular nucleus of the thalamus | RT |  |
| 1085 | Lateral habenula | LH |  |
| 1086 | Medial habenula | MH |  |
| 1087 | Hypothalamus | HY |  |
| 1088 | Medial preoptic nucleus | MPN |  |
| 1089 | Anterior hypothalamic nucleus | AHN |  |
| 1090 | Ventromedial hypothalamic nucleus | VMH |  |
| 1091 | Posterior hypothalamic nucleus | PH |  |
| 1092 | Mammillary body | MBO |  |
| 1093 | Suprachiasmatic nucleus | SCH |  |
| 1094 | Arcuate hypothalamic nucleus | ARH |  |
| 1095 | Paraventricular hypothalamic nucleus | PVH |  |
| 1096 | Lateral preoptic area | LPO |  |
| 1097 | Tuberal nucleus | TU |  |
| 1098 | Lateral hypothalamic area | LHA |  |
| 1099 | Subthalamic nucleus | STN |  |
| 1100 | Zona incerta | ZI |  |
| 1101 | Superior colliculus, sensory related | SCs |  |
| 1102 | Superior colliculus, motor related | SCm |  |
| 1103 | Inferior colliculus | IC |  |
| 1104 | Inferior colliculus, central nucleus | ICc |  |
| 1105 | Parabigeminal nucleus | PBG |  |
| 1106 | Pedunculopontine nucleus | PPN |  |
| 1107 | Pretectal region | PRT |  |
| 1108 | Oculomotor region | IIIR | III, EW |
| 1109 | Periaqueductal gray | PAG |  |
| 1110 | Red nucleus | RN |  |
| 1111 | Midbrain reticular nucleus | MRN |  |
| 1112 | Cuneiform nucleus | CUN |  |
| 1113 | Interpeduncular nucleus | IPN |  |
| 1114 | Ventral tegmental area | VTA |  |
| 1115 | Substantia nigra, compact part | SNc |  |
| 1116 | Substantia nigra, reticular part | SNr |  |
| 1117 | Dorsal nucleus raphe | DR |  |
| 1118 | Pons | P |  |
| 1119 | Nucleus of the lateral lemniscus | NLL |  |
| 1120 | Superior olivary complex | SOC |  |
| 1121 | Parabrachial nucleus | PB |  |
| 1122 | Principal sensory nucleus of the trigeminal | PSV |  |
| 1123 | Pontine central gray | PCG |  |
| 1124 | Pontine gray | PG |  |
| 1125 | Trigeminal motor complex | VMc | V, P5, I5, Acs5 |
| 1126 | Facial motor nucleus | VII |  |
| 1127 | Dorsal tegmental nucleus | DTN |  |
| 1128 | Laterodorsal tegmental nucleus | LDT |  |
| 1129 | Tegmental reticular nucleus | TRN |  |
| 1130 | Pontine reticular formation | PRF | PRNc, PRNr |
| 1131 | Raphe ventral complex | RVC | IF, IPN, RL, CLI, CS |
| 1132 | Hemispheric regions | HEM |  |
| 1133 | Vermal regions | VERM |  |
| 1134 | Paraflocculus | PFL |  |
| 1135 | Floccular nodular region | FLNOD | NOD, FL, UVU |
| 1136 | Dentate nucleus | DN |  |
| 1137 | Interposed nucleus | IP |  |
| 1138 | Fastigial nucleus | FN |  |
| 1139 | Vestibulocerebellar nucleus | VeCB |  |
| 1140 | Nucleus of the trapezoid body | NTB |  |
| 1141 | Spinal nucleus of the trigeminal region | SPVr | SPVC, SPVI, SPVO |
| 1142 | Nucleus of the solitary tract | NTS |  |
| 1143 | Dorsal cochlear nucleus | DCO |  |
| 1144 | Ventral cochlear nucleus | VCO |  |
| 1145 | Vestibular nuclei | VNC |  |
| 1146 | Spinal vestibular nucleus | SPIV |  |
| 1147 | Dorsal column nuclei | DCN |  |
| 1148 | Inferior olivary complex | IO |  |
| 1149 | Perihypoglossal nuclei | PHY |  |
| 1150 | Dorsal motor nucleus of the vagus nerve | DMX |  |
| 1151 | Hypoglossal nucleus | XII |  |
| 1152 | Medullary reticular zone | MRz | GRN, LRN, MARN, MDRN, PARN, PGRN, |
| 1153 | Intermediate reticular nucleus | IRN |  |
| 1154 | Medulla, raphe, behavioral state related | MY-sat |  |
| 1155 | fiber tracts | fiber tracts |  |
| 1156 | corpus callosum | cc | ccb, ccg, ccs, fp |
| 1157 | cingulum bundle | cing |  |
| 1158 | anterior commissure | ac | act, aco |
| 1159 | optic tract and chiasm | optc | opt,och |
| 1160 | dorsal hippocampal commissure | dhc |  |
| 1161 | fimbria | fi |  |
| 1162 | columns of the fornix | fx |  |
| 1163 | stria terminalis | st |  |
| 1164 | thalamus related | lfbst |  |
| 1165 | medial forebrain bundle | mfb |  |
| 1166 | fasciculus retroflexus | fr |  |
| 1167 | posterior commissure | pc |  |
| 1168 | corticospinal tract | cst |  |
| 1169 | medial lemniscus | ml |  |
| 1170 | lateral lemniscus | ll |  |
| 1171 | medial longitudinal fascicle | mlf |  |
| 1172 | superior cerebelar peduncles | scp |  |
| 1173 | inferior cerebellar peduncle | icp |  |
| 1174 | trigeminal nerve | Vn |  |
| 1175 | cochlear nerve | cVIIIn |  |
| 1176 | lateral ventricle | VL |  |
| 1177 | third ventricle | V3 |  |
| 1178 | fourth ventricle | V4 |  |
| 1179 | cerebral aqueduct | AQ |  |
| 1180 | Whole brain _root_ | Brain |  |

**Table S2** Regions of interest for left hemisphere (ROI 1-180) and right hemisphere (ROI 1001-1180) for reduced CCFv3 (i.e. r1CCFv3 label set).

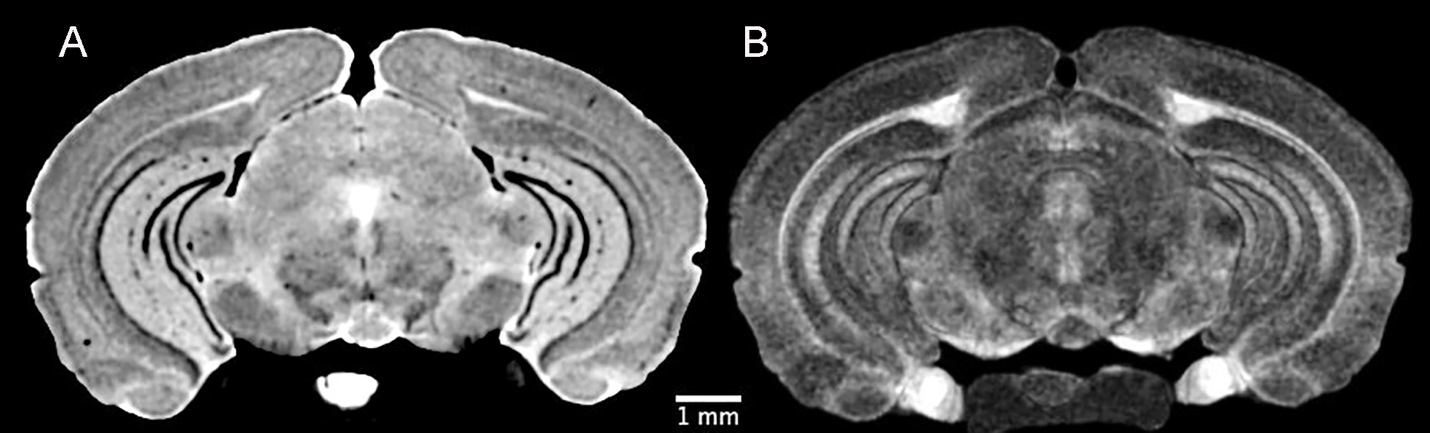

**Figure S3** Representative A)DWI and B) QA images from higher throughput (42 hr) 25 μm, 61 angle acquisition

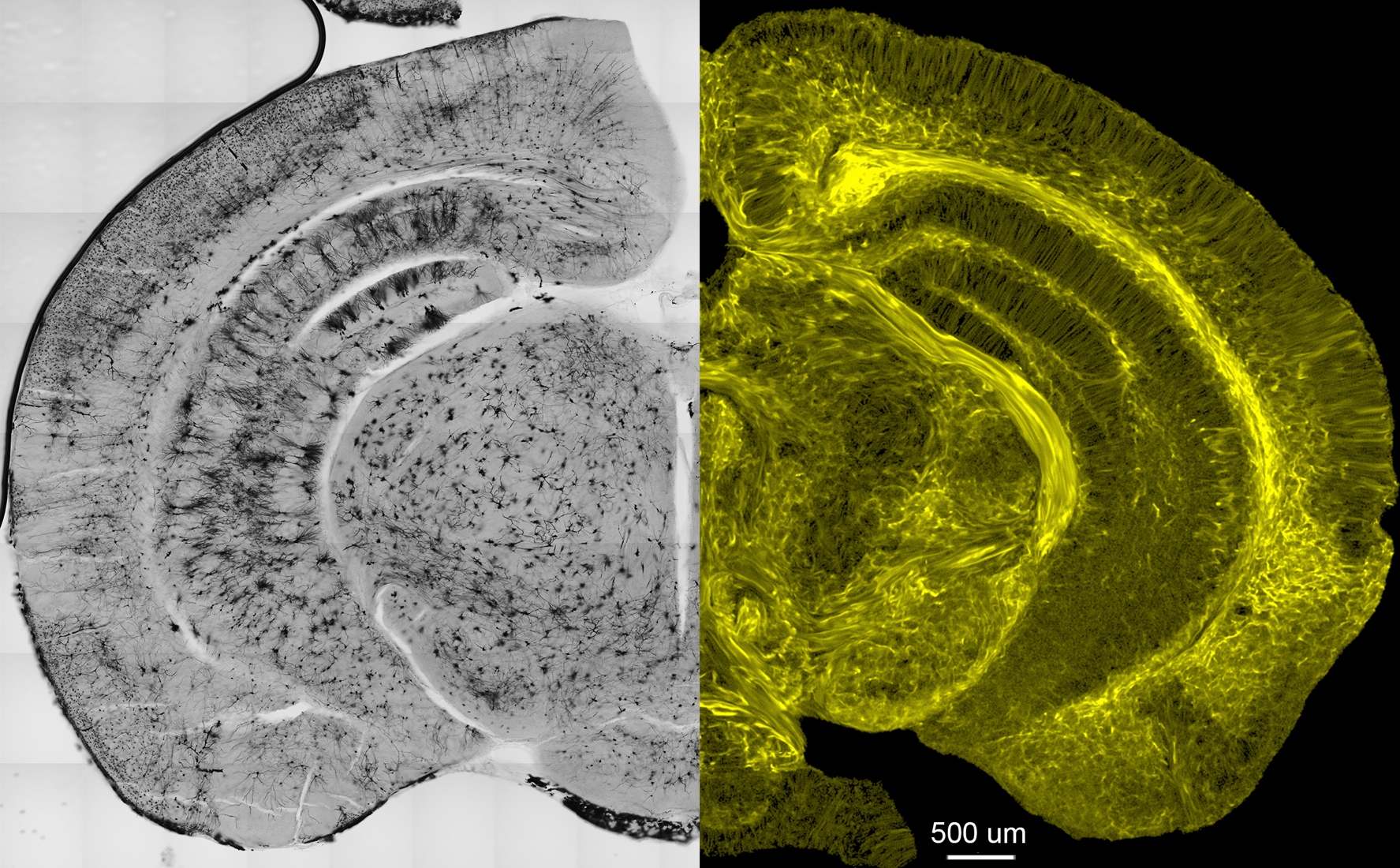

**Figure S4** Comparison of A) Golgi stain and B) Super resolution TDI @ 5 mm resolution (Specimen 190415-2:1)

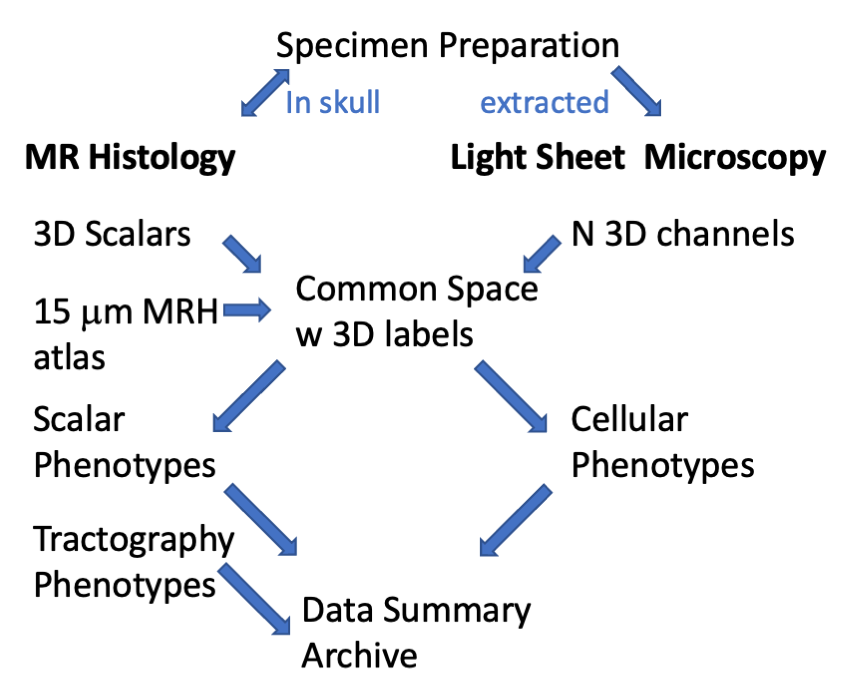

**Figure S5** Workflow for acquisition of combined whole brain MR histology and light sheet microscopy. The actively stained brain (still in the cranium) is scanned in a 9.4T MR imaging system equipped with bespoke coils achieving gradients more than 100 times those of a clinical scanner. Compressed sensing is used to accelerate a high angular diffusion tensor acquisition. A series of image pipelines process the large (300 GB) 4D MRI data to derive scalar and tractography data. A new MR atlas acquired at 15 μm isotropic resolution includes a subset of 360 labels from the Allen Brain Atlas Common Coordinate Framework. A second pipeline maps these labels onto the strain under study. The brain is removed from the skull, cleared (SHIELD), stained (SWITCH) and scanned @ 1.8 X1.8 x 4 μm. A third pipeline registers the light sheet data to the common space defined by the MR histology image of the strain under study (in the skull) removing the distortions that accompany LSM. Image derived phenotypes from the 360 ROI are aggregated into a final summary.

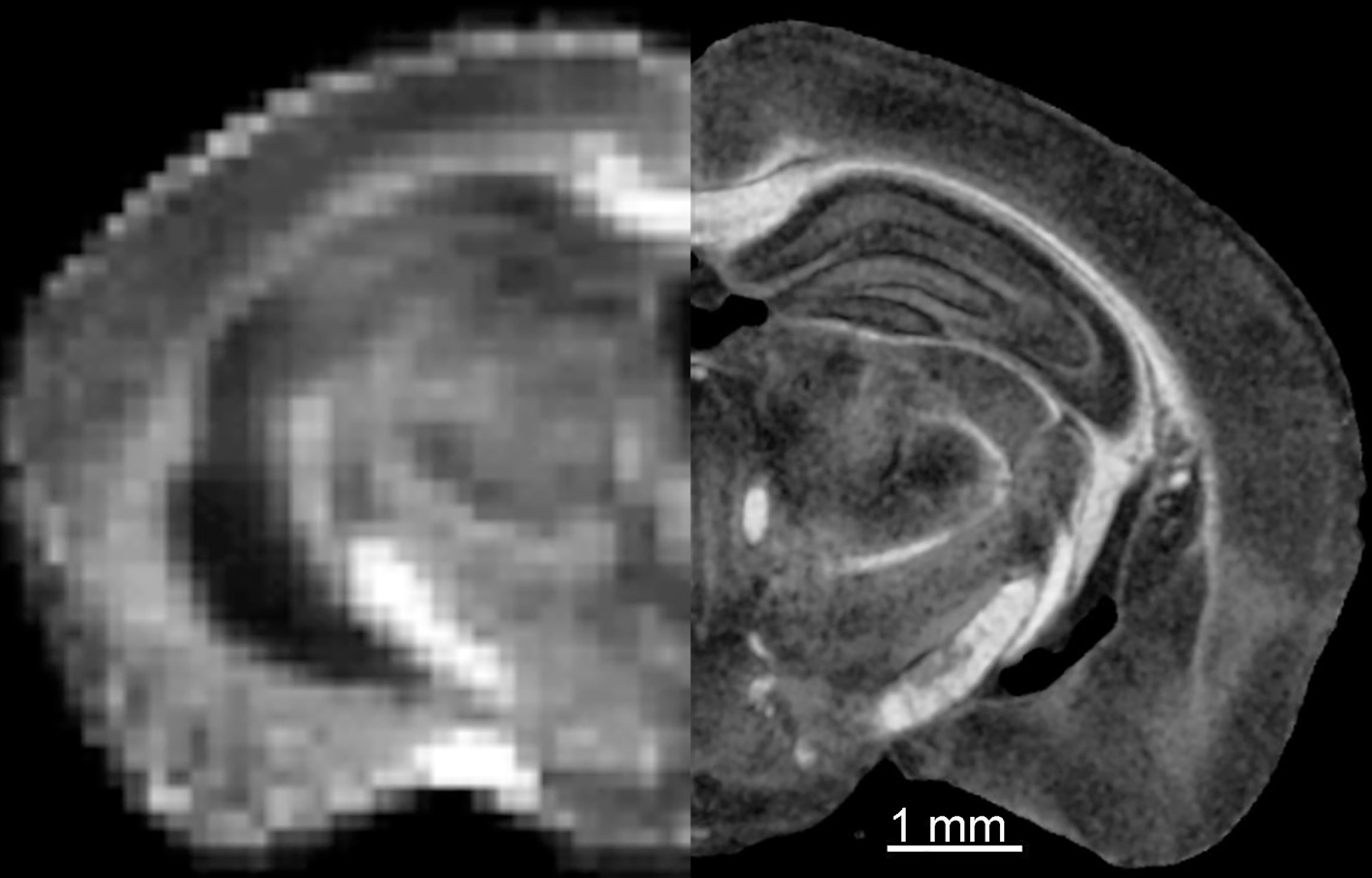

**Figure S6** Comparison in vivo fractional anisotropy image @ 150 μm^3^ (left) with ex vivo MR histology fractional anisotropy image @ 15 μm^3^ (right). In vivo image Courtesy of Professor Ian Shih, University of North Carolina (https://www.med.unc.edu/bric/camri/imaging-service/mouse-brain-in-vivo-epi-dti/)

**Figure S7** High performance computer infrastructure to facilitate combined MRH and LSM with large multidimensional images. Key is the division of the workflow into 5 stages—acquisition, reconstruction, archive, post-processing, and sharing with computational resources optimized for large data files at each stage.

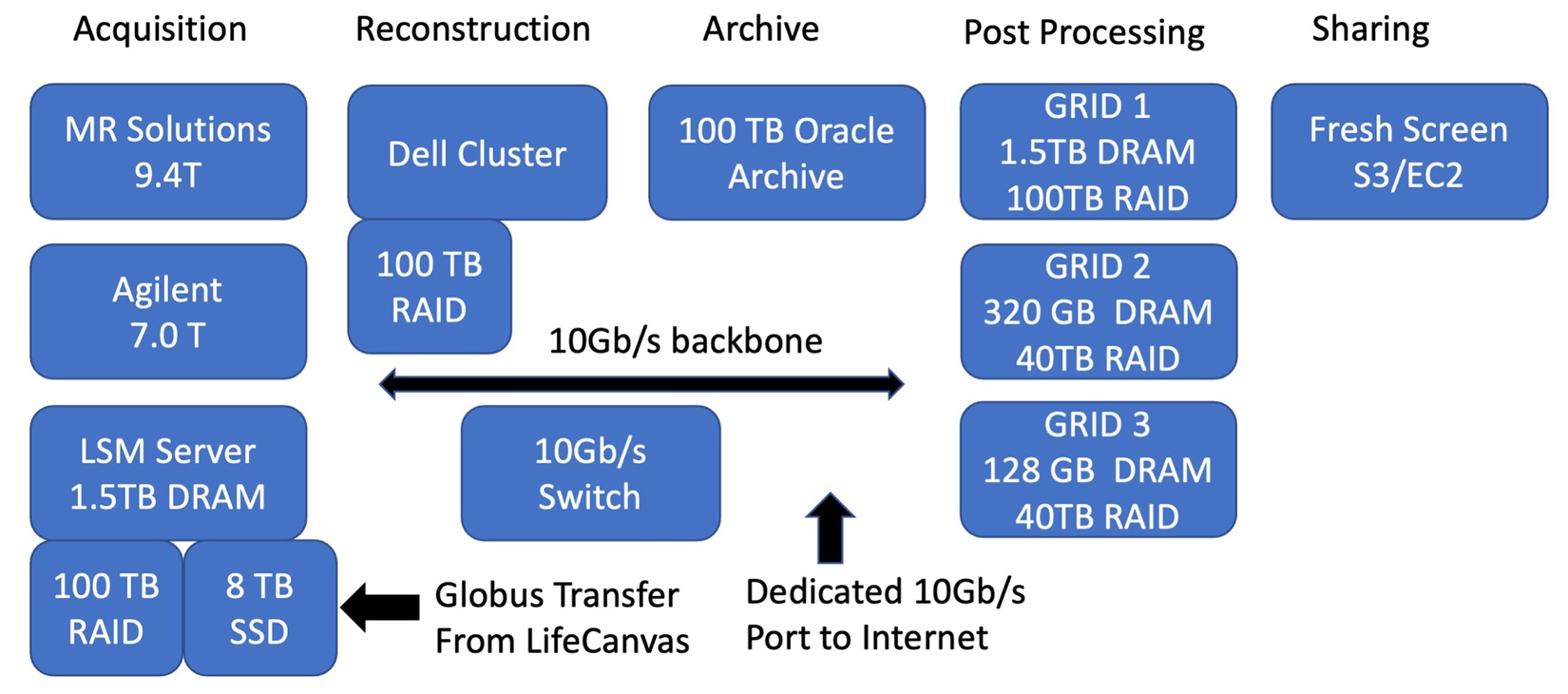

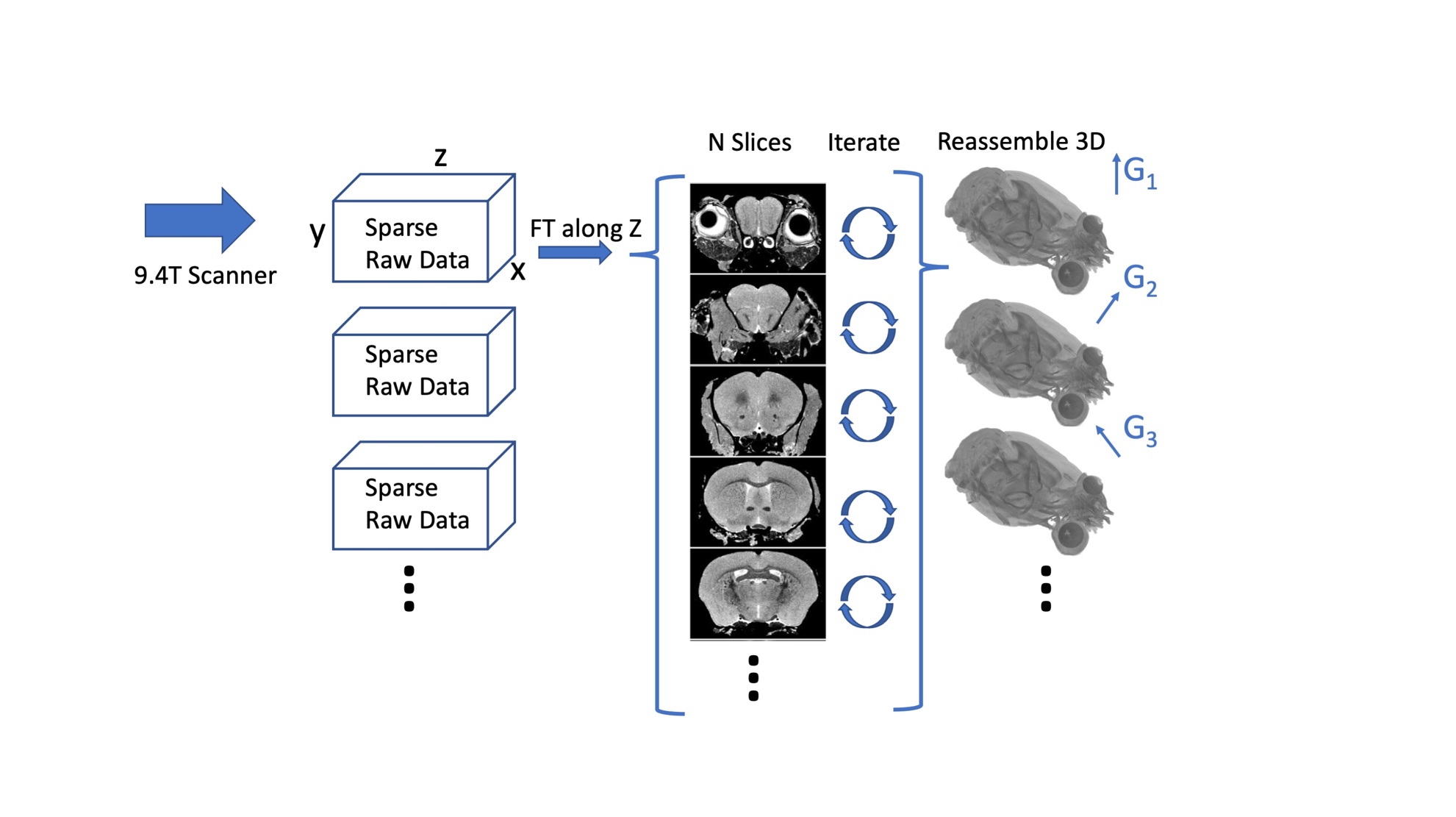

**Figure S8.** A pipeline integrates a 4-dimensional acquisition from the scanner with compressed sensing reconstruction in a high performance Dell cluster. Data from the scanner is streamed from the scanner to the cluster. Fourier space is under sampled by a factor of 8 using a probabilistic distribution along two (phase) dimensions of acquisition. A Fourier transform along the fully sampled readout axis produces up to 2000 2 dimensional arrays which are launched as individual iterative reconstructions in the cluster. Upon completion of the iterative reconstruction, these 2D arrays are reassembled into a 3D volume. A script on the scanner launches the

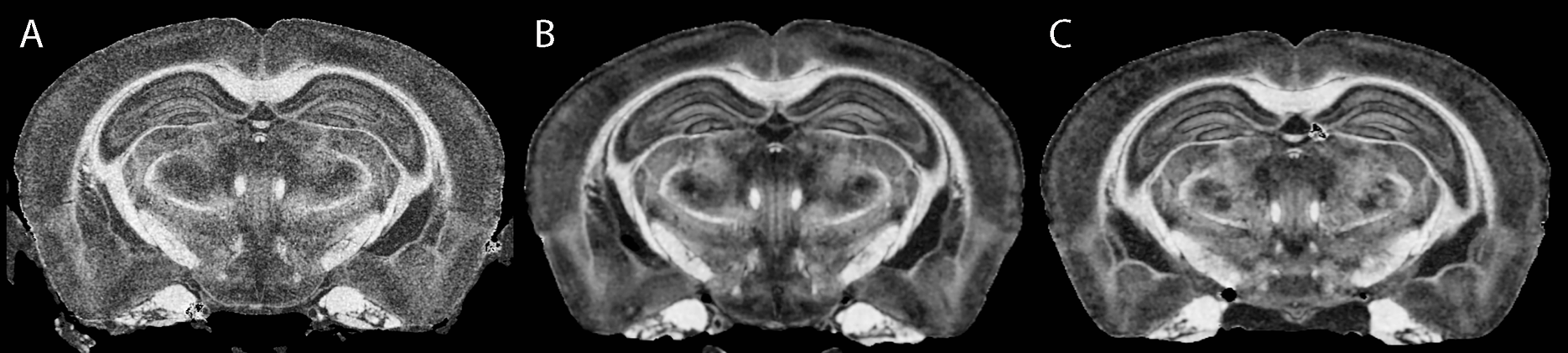

Figure S9 Comparison of spatial and angular resolution A) Specimen 190415-2:1: 15 μm, 61 angles; B) 190415-1:1 25 μm, 126 angles; C) 25 μm, 61 angles

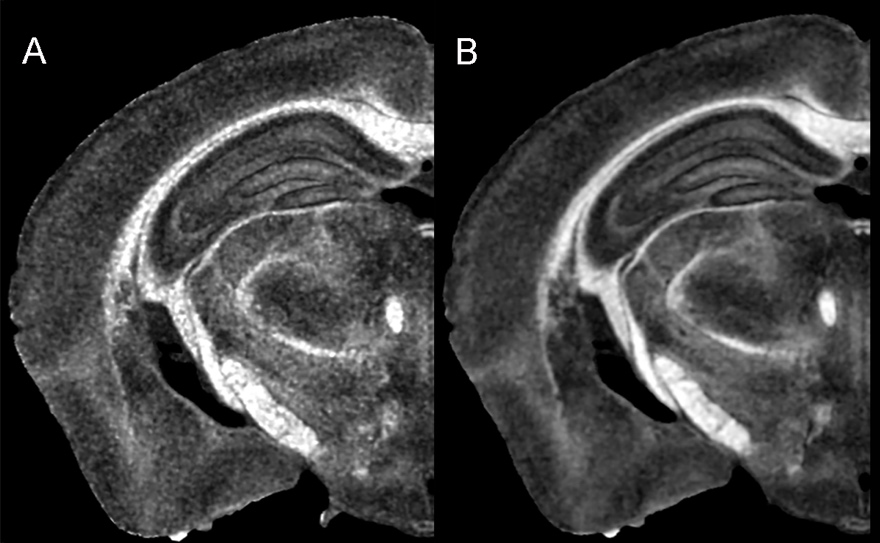

**Figure S10** Specimen 200316-1:1(Table 1) @ 15 μm, isotopic spatial resolution A) before denoising; B) after denoising
